# Human TUBA1B short open reading frame product regulates cancer cell growth via importin β

**DOI:** 10.1101/2023.08.26.554759

**Authors:** Yaling Tao, Xuefeng Bai, Yinjie Zhou, Yue Zhao, Liangwei Yang, Shun Zhang, Huina Liu, Xiaochun Huang, Edoardo Schneider, Anna Zampetaki, Andriana Margariti, Mauro Giacca, James N. Arnold, Lingfang Zeng, Ting Cai

## Abstract

Understanding cancer biology is crucial for improving treatment strategies. This study identified TUBA1B-sORF1, a short open reading frame product alternatively translated from the human α-tubulin gene (TUBA1B), which has a completely different amino acid sequence from the α-tubulin 1B chain. TUBA1B-sORF1 is highly expressed in cancer cell lines and gastric carcinoma. Both methionine-initiated canonical and leucine-initiated noncanonical translations of TUBA1B-sORF1 coexist in cancer cells, and there is a transition between sORF1 and α-tubulin translations, evidenced by the TUBA1B-sORF1^⁺^/α-tubulin^low/-^ subpopulation. Knocking down TUBA1B-sORF1 reduces cancer cell proliferation and tumorigenicity. TUBA1B-sORF1 facilitates protein nuclear translocation, leading to the upregulation of proliferation-promoting genes and downregulation of proliferation-inhibiting genes. Specifically, it forms a complex with importin β and β-catenin, promoting β-catenin nuclear translocation and target gene transcription. These findings reveal that TUBA1B is a polycistronic gene translating at least two entirely different proteins: α-tubulin and TUBA1B-sORF1. The variable translation between them may regulate tumorigenesis, making TUBA1B-sORF1 a promising therapeutic target and diagnostic biomarker for cancer treatment.

## Introduction

Understanding the complex mechanisms driving cancer progression is crucial for developing innovative therapeutic strategies that can effectively combat tumour recurrence and metastasis (Sung, 2021). Chemotherapy has long been established as a cornerstone of cancer treatment, aiming to eliminate tumour cells through various mechanisms, including cell cycle arrest, DNA damage, and disruption of cellular structures critical for survival (Mitchison, 2012). Microtubules are hollow cylindrical tubes made of heterodimers of α-tubulin and β-tubulin in the cytoplasm and play crucial roles in cell morphology, migration, intracellular transport, organelle stability and cell division. A dynamic transition exists between polymerization and depolymerization via the addition and subtraction of tubulin units, respectively, at both ends of the microtubule structure, depending on cellular function requirements (Wilson & Jordan, 1995). The microtubule dynamic transition is more active in cancer cells than in normal cells. Among chemotherapies, microtubule-targeting agents (MTAs) have emerged as a prominent therapeutic approach due to their ability to disrupt the dynamic transition either by inhibiting the addition of tubulin subunits such as Vinca alkaloids or preventing depolymerization such as taxane, consequently suppressing cell mitosis and eventually leading to cell death (Dumontet & Jordan, 2010; Wordeman & Vicente, 2021). MTAs have been used as a first-line chemotherapy for breast, lung, and ovarian cancer (Čermák, 2020). The response to MTA therapy is generally favourable. However, cancer recurrence frequently occurs after MTA treatment. Numerous studies have aimed to explain the mechanisms behind the recurrence phenomenon, and different theories have been proposed (Smith *et al*, 2022; Kavallaris, 2010; Dan *et al*, 2021). However, the underlying mechanisms remain unsolved.

Recent high-throughput sequencing studies have reported abnormal levels of the α-tubulin 1b chain (TUBA1B) in cancer, establishing it as a valuable prognostic and overall survival indicator. Hu *et al*. reported a strong correlation between elevated mRNA expression levels and overall survival in patients with colorectal cancer (Hu, 2021). Similarly, Xiong *et al*. reported significantly increased TUBA1B mRNA levels in hepatocellular carcinoma (Xiong, 2019). In pancreatic ductal adenocarcinoma, Qin *et al*. constructed a classifier that identified TUBA1B mRNA as a promising biomarker for patient prediction (Qin, 2021). Despite its importance in maintaining normal cell function, the underlying mechanism and function of the α-tubulin protein in cancer remain elusive. Intriguingly, Linge *et al*. demonstrated reduced TUBA1B protein levels in uveal melanomas that metastasized compared to those that did not, indicating the intricate and multifaceted involvement of TUBA1B in cancer progression (Linge, 2012). The discrepancy between the mRNA and protein levels implies that other mechanisms are involved in protein translation from mRNAs.

In the last two decades, great progress has been made in understanding protein translation, including non-AUG codon initiation of translation, stop-codon readthrough, overlapping open reading frames, short open reading frames (sORFs), and functional ORFs in so-called noncoding RNAs, etc (Adhikari, 2020; Wright *et al*, 2022; Cao, 2021; Vanderperre, 2011; Bergeron, 2013; Chalick, 2016). These events dramatically increase the cell proteome and are possibly involved in almost all physiological/pathophysiological processes, including those exploited in cancer. sORFs, defined as a subcategory of ORFs consisting of sequences shorter than 300 nucleotides or 100 codons, have traditionally been overlooked or excluded from datasets due to a lack of coding potential. In recent years, there has been a marked increase in the identification of sORFs with the capacity to encode proteins, thus driving the databases collecting information on sORFs (Guerra-Almeida & Nunes-da-Fonseca, 2020; Brunet, 2021). In this study, we found a novel sORF product overlapping with the human gene TUBA1B with 50 codons, named TUBA1B-sORF1 or sORF1.

## Results

### The short open reading frame product is translated in the *TUBA1B* gene

The aforementioned studies indicate that the TUBA1B gene may play a significant role in cancer and that discrepancies exist between its mRNA and protein levels. Indeed, dramatic variation existed in α-tubulin but not in β-tubulin among different cancer tissues, as revealed by Western blot analysis (Figure 1A). The variation of a protein level can be derived from the regulation of either translation from its mRNA or protein stability. We hypothesized that the variation in α-tubulin within different cancer tissues was derived from the translation shift between the main ORF (mORF) for α-tubulin and the short ORF(s) and that the sORF product(s) played a critical role in cancer cell cellular functions.

**Figure 1.**
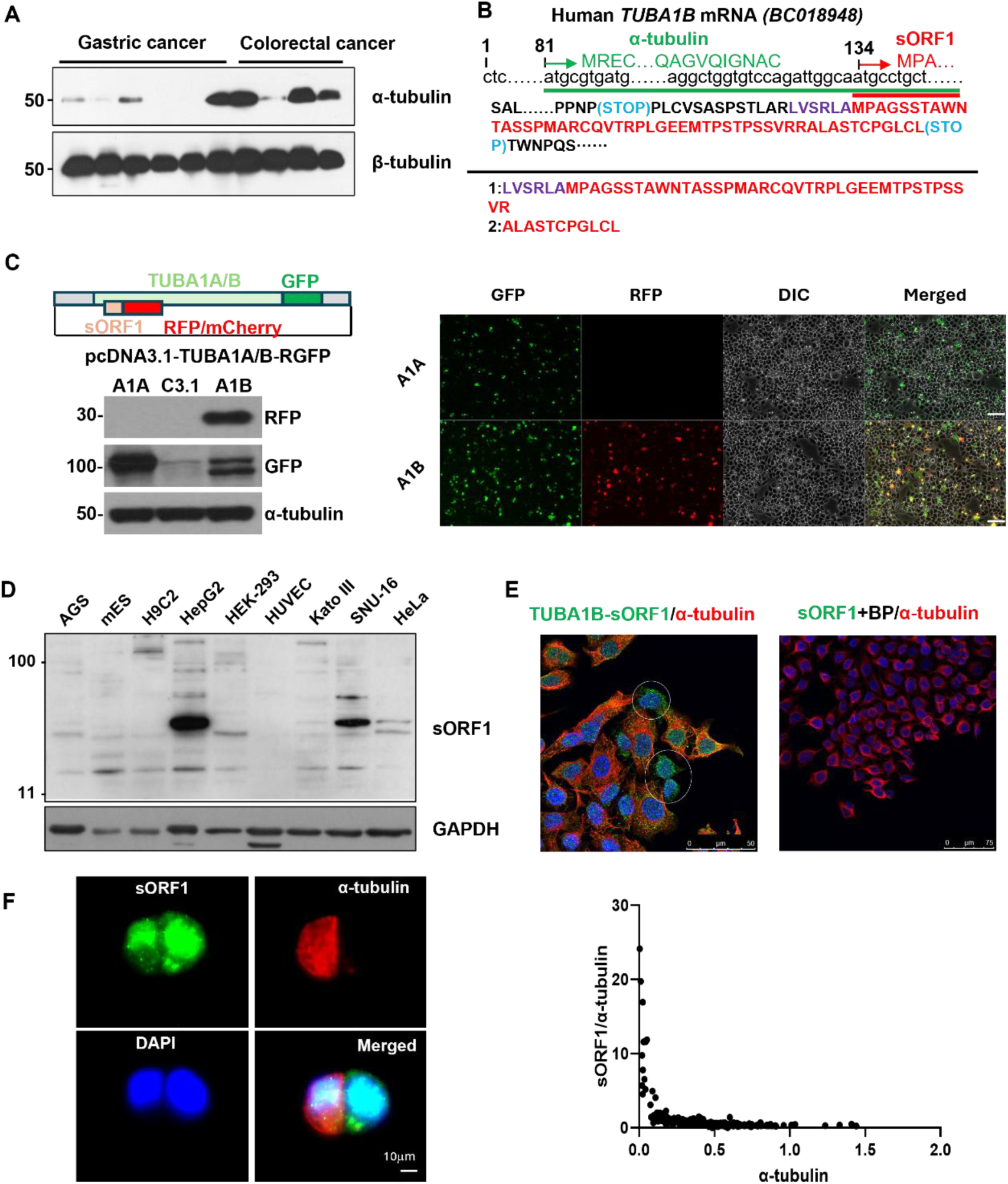
A short open reading frame product was alternatively translated from the human *TUBA1*B gene. (A) Western blot analysis, performed on equal amounts of samples in the same batch, revealed varying levels of α-tubulin and stable levels of β-tubulin in gastric and colorectal cancer tissues. (B) Schematic representation illustrating the main ORF (a-tubulin in green) and sORF1 (in red) in TUBA1B mRNA (upper panel), along with two peptide fragments identified through proteomic analysis of HeLa and SNU-16 cell lysates (lower panel). (C) Transfection of HeLa cells with *pcDNA3.1-TUBA1A-RGFP* (*A1A*) or *pcDNA3.1-TUBA1B-RGFP* (*A1B*) plasmids and subsequent WESTERN BLOT analysis (left) with the indicated antibodies and observation under an inverted fluorescence microscope (right). pcDNA3.1 was included as a mock control (C3.1). The mCherry signal represents sORF1, while the GFP signal signifies α-tubulin. Scale bar: 100 μm. (**D**) Identification of TUBA1B-sORF1 in multiple cancer cell lines through blotting with a peptide-generated antibody. AGS: human gastric cancer, mES: mouse embryonic stem cells, H9C2: rat cardiomyocytes, HepG2: human hepatocarcinoma, HEK-293: human embryonic kidney epithelial cells, HUVEC: human umbilical vein endothelial cells, Kato III: human gastric cancer, SNU-16: human gastric cancer, and HeLa: human cervical cancer. (**E**) Immunofluorescence staining was performed in HeLa cells with anti-sORF1 (green) and anti-α-tubulin (red) antibodies. The white circle shows sORF1^+^/α-tubulin^-^ cells. BP: peptide blocking. (**F**) The translation of sORF1 and α-tubulin was negatively related in SNU-16 cells, as revealed by immunofluorescence staining with anti-sORF1 (green) and anti-α-tubulin (red) and fluorescence intensity analysis of 200 stained SNU-16 cells (right panel).

To test this hypothesis, the translation potency of both the human and mouse *TUBA* genes was first analysed. Highly conserved sORF1 exists in all isoforms of both human and mouse *TUBA* genes (Figure S1). To assess whether this sORF1 was encodable, proteins with molecular weights between 3 kDa and 25 kDa from the cervical cancer cell line HeLa and gastric cancer cell line SNU-16 were subjected to proteomic analysis to search for peptide fragments from the sORF1 sequences of different isoforms. Two peptide fragments were detected, and upon concatenation, they accurately covered the full sORF1 sequence within the *TUBA1B* gene but not within other isoforms (Figure 1B and S2). An additional six amino acids (LVSRLA) upstream of the start codon methionine were detected in peptide 1 (Figure 1B), implying upstream noncanonical non-AUG initiation of *TUBA1B-sORF1*. Peptide 1 is highly conserved in the *TUBA1A* gene, except for one amino acid. The start codon (AUG) of *TUBA1A-sORF1* is located far upstream of the peptide 1-like sequence. The putative canonical translation of *TUBA1A-sORF1* will yield a 134-amino acid product, the trypsin digestion of which can produce the peptide 1-like sequence. In this case, noncanonical non-AUG initiation can be excluded.

To determine whether *TUBA1B* mRNA was the sole source of this sORF1 product, the full-length *TUBA1A* (transcript variant 1, *AB590174.1*) and *TUBA1B* (*K00558.1*) cDNAs were subsequently cloned and inserted into the pcDNA3.1+ vector, in which mCherry (red fluorescence protein, RFP) and green fluorescence protein (GFP) were inserted in-frame upstream of the stop codon of sORF1 and mORF, respectively, which were designated as *pcDNA3.1-TUBA1A-RGFP* (*A1A*) and *pcDNA3.1-TUBA1B-RGFP* (*A1B*) (Figure 1C upper left). Transfection of these plasmids into HeLa cells detected GFP signals from both plasmids but detected RFP signals only in *A1B* cells, as revealed by Western blot (Figure 1C, lower left) and inverted fluorescence microscopy (Figure 1C, right). Intriguingly, the intensity of RFP and GFP in *A1B*-transfected cells displayed an inversely proportional relationship, implying that a translational shift may exist between sORF1 and mORF in different cells.

To further assess endogenous TUBA1B-sORF1 expression, polyclonal antibodies were generated from rabbits by immunopeptides (immunogen: QVTRPLGEEMTPSTC). Methionine-initiated sORF1 is expected to contain only 50 amino acids, corresponding to a product of approximately 5-6 kDa. However, Western blot analysis of various cell lysates revealed multiple bands with different molecular weights for endogenous sORF1, with the main band appearing at approximately 50 kDa, while no bands were detected below 10 kDa (Figure 1D). The higher bands may be due to some type of posttranslational modification (PTM). Moreover, the expression level of sORF1 was higher in cancer cell lines than in primary or immortalized normal cells. To confirm the specificity of these bands, an overnight antibody blocking with immunopeptide was performed. As expected, no bands were detected after blocking (Figure S3), indicating the specificity of the peptide-generated antibody. Subsequently, immunofluorescence (IF) staining with unblocked and blocked antibodies in HeLa cells revealed the presence of specific sORF1 signals, while there was no colocalization with α-tubulin (Figure 1E). Similarly, IF staining of HeLa and SNU-16 (gastric carcinoma cell line) cells revealed uneven expression levels of the internal sORF1 product and α-tubulin, with a subpopulation showing sORF1^+^/α-tubulin^low/-^ (Figure 1E, white circle and 1F, left). These cells accounted for approximately 5-10% of the total population (Figure S4), and intensity analysis showed that the ratio of sORF1/α-tubulin had an inversely proportional relationship (Figure 1F, right).

These results confirm the sole source of sORF1 from the *TUBA1B* gene and the competitive translation selection between the mORF and sORF. The variable translation between mORF and sORF may reflect a unique cell type or status in the cancer cell population.

### The non-AUG initiation of *TUBA1B-sORF1* can occur from different codons

As described above, the LVSRLA sequence in peptide 1 (Figure 1B) implies non-AUG codon initiation within *TUBA1B-sORF1*. To confirm this, two strategies involving artificial plasmid construction and CRISPR/Cas9 homologous DNA recombination-mediated gene editing were adopted to insert a FLAG tag in-frame upstream of the first methionine codon of *TUBA1B-sORF1*. First, the *pSh2-TUBA1B-FsORF1* plasmid was created (Figure 2A, upper). Co-immunoprecipitation (Co-IP) was performed in transfected HeLa cells, followed by Western blot with anti-sORF1 and anti-FLAG antibodies with the FLAG-positive plasmid (*pSh2-F-HD3*, FLAG-tagged HDAC3), mock vector *pSh2* and TE as vehicle controls. As shown in Figure 2A lower, the same ∼50 kDa band positive for both sORF1 and FLAG was detected. Second, CRISPR/Cas9-mediated gene editing with a guide RNA (GTGACCACCTAGTCTAAAGT, located in intron 1) and a homologous recombination template (*pUC57-TUBA1B-FsORF1*) was performed to endogenously insert the FLAG tag sequence in-frame upstream of *TUBA1B-sORF1* in SNU-16 cells (Figure 2B, upper). DNA gel electrophoresis of polymerase chain reaction (PCR) product showed successful gene editing (Figure 2B, lower). Multiple positive FLAG bands were detected by Western blot, where the smallest band was lower than 11 kDa, matching the putative FLAG-sORF1 product, while more bands appeared as the molecular weight increased to 180 kDa (Figure 2C, red arrows). IF staining also revealed merged sORF1 and FLAG signals within the same cell (Figure 2D). These data directly confirmed the non-AUG codon initiation of *TUBA1B-sORF1*.

**Figure 2.**
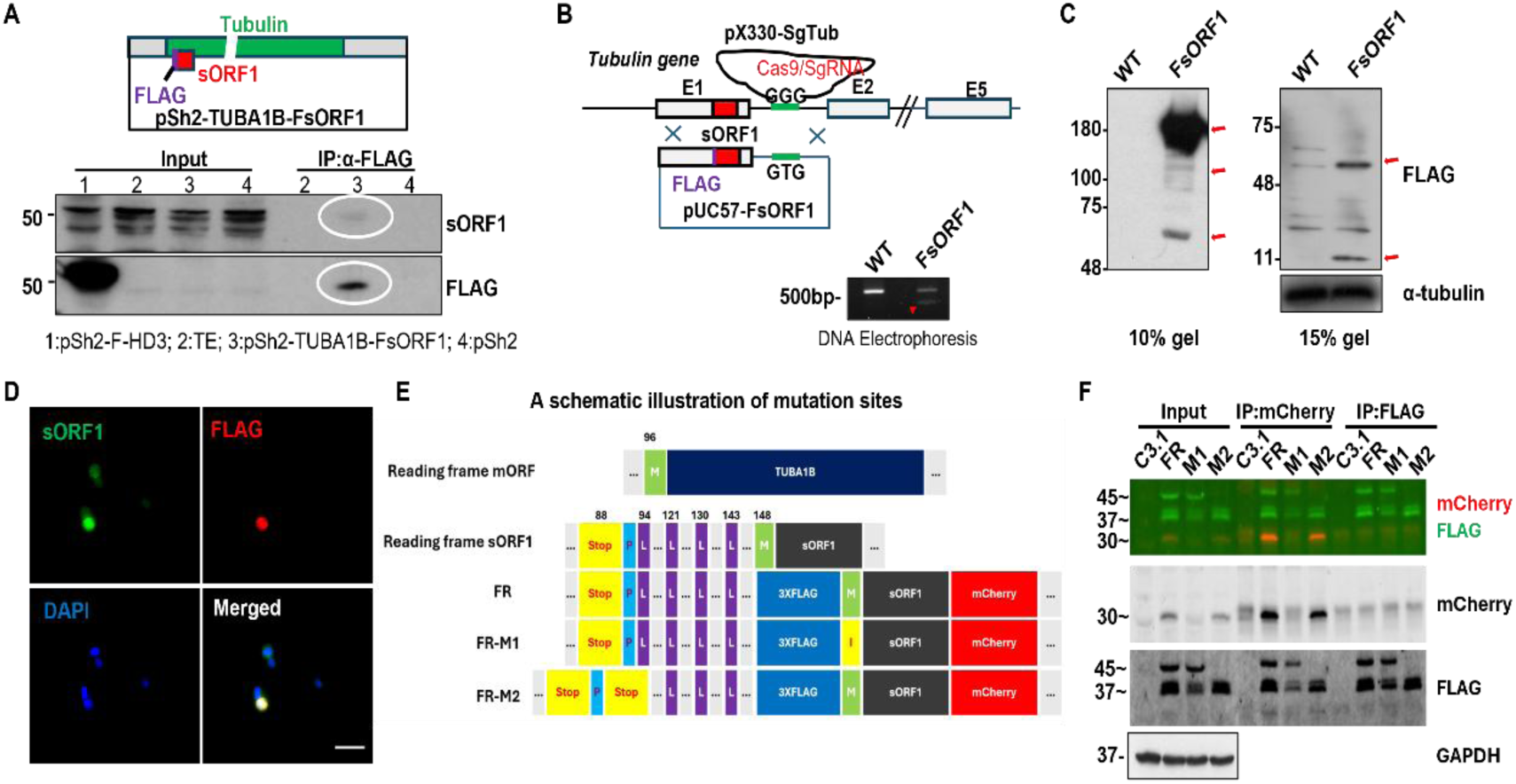
Non-AUG codon initiation occurred during TUBA1B-sORF1 translation. (**A**) HeLa cells were transfected with pSh2-TUBA1B-FsORF1 with a FLAG tag inserted upstream of the AUG codon of sORF1 (upper panel), followed by immunoprecipitation with anti-FLAG and immunoblotting with anti-sORF1 and anti-FLAG antibodies. The white circle shows the FLAG-tagged sORF1 band translated from noncanonical initiation. (**B-D**) CRISPR/Cas9 plus homologous recombination strategy (**B**) was used to insert a FLAG tag upstream of the AUG codon in the TUBA1B-sORF1 in the SNU-16 cell line, followed by verification via PCR with primers targeting the FLAG sequence (**C**, upper left) and WESTERN BLOT with anti-FLAG (C, lower left and right) and immunofluorescence staining (**D**) with anti-FLAG (red) and anti-sORF1 (green) with DAPI included to counterstain nuclei. Scale bar: 100μM. (**E**) A schematic figure illustrating the verification of the TUBA1B-sORF1 initiation codon. The 3XFLAG tag was inserted upstream of the methionine, while mCherry was added to the C-terminus of sORF1 as the FR plasmid. Based on the FR, all the methionine codons located within the sORF1 sequence were mutated to isoleucine, as the FR-M1 plasmid. The leucine at position 94 was mutated to a stop codon as in the plasmid FR-M2. (**F**) HeLa cells were transfected with FR, FR-M1 or FR-M2, followed by IP and WESTERN BLOT with the indicated antibodies. Fluorescein-conjugated secondary antibodies were used to visualize the bands.

An increasing number of studies have revealed that noncanonical translation is often initiated by leucine (Sanz et al., 2019; Schwab et al., 2004; Starck, 2012). The detection of LVSRLA in peptide 1 upstream of the first methionine of sORF1 implies CUG codon initiation. However, an arginine residue is located upstream of the leucine residue, suggesting that peptide 1 can be a trypsin cleavage product from the initiation of another non-AUG codon in the 5’ undetermined region. Analysis of the *TUBA1B* mRNA sequence (NM_006082) revealed four leucine residues (at nucleotides 94, 121, 130 and 143) positioned between a stop codon (88) and the methionine (148) within the *TUBA1B-sORF1 frame*, where the first leucine codon (94) is only two nucleotides away from the methionine (96) of α-tubulin (Figure 2E). Additionally, the results from the GWIPS-viz Genome Browser (Kiniry et al., 2018) for *TUBA1B* did not detect any sequences representing the methionine of sORF1 or the preceding three leucine residues. Thus, we hypothesize that the first leucine (94, **<u>cta</u>**tgc) can initiate translation of TUBA1B-sORF1. To determine this, based on the full-length mRNA of *TUBA1B*, a 3XFLAG tag and an mCherry tag were inserted upstream of the methionine codon and the stop codon of sORF1, respectively. The resulting plasmid was designated *pcDNA3.1-TUBA1B-FLAG-sORF1-mCherry* (abbreviated as *FR*). Subsequently, all the methionine codons of sORF1 were mutated to isoleucine (M^AUG^/I^AUC^) to obtain FR-M1. Separately, leucine (94) was mutated to a stop codon to obtain FR-M2 (94, **<u>taa</u>**tgc, Figure 2E). After transfection into HeLa cells, multiple bands were detected via Co-IP and Western blot with anti-mCherry and anti-FLAG antibodies (Figure 2F). The ∼30 kDa band was positive for mCherry which matched the molecular weight of the sORF1-mCherry fusion protein, but negative for FLAG. The ablation of this band by the M^AUG^/I^AUC^ mutation indicates that the translation of sORF1 can be initiated from the first methionine site (FR-M1 in Figure 2F), which is possibly the main product. There were three bands positive for FLAG (one band at ∼45 kDa and two at ∼37 kDa), which were also positive for mCherry, as expected, suggesting that three upstream non-AUG codons can initiate the translation of sORF1. The ∼45 kDa band disappeared when the CUA codon was mutated to a stop codon (UAA) (FR-M2 in Figure 2F), indicating initiation from the leucine codon (94, CUA). The initiation sites of the other two FLAG-positive bands (∼37 kDa) need to be characterized via further investigation to determine whether they are from sORF1 non-AUG codons or from FLAG itself.

There could be post-translational modifications (PTMs) in certain amino acid residues between the codons for leucine (94, CUA) and methionine (148), as the discrepancy in residue numbers did not align with the difference in molecular weight. Despite there being only 9 amino acids, the molecular weight difference is approximately 10 kDa, indicating additional mass unaccounted for. Taken together, our data demonstrated that both methionine (148) and one or more upstream non-methionine codon(s), especially the leucine (94) codon, can initiate the translation of TUBA1B-sORF1. There may exist a selection between leucine (94) for sORF1, and methionine (96, ct**<u>atg</u>**c) for α-tubulin in some cells, leading to the occurrence of the sORF1^+^/α-tubulin^low/-^ subpopulation in cancer cells.

### sORF1 expression is essential for cancer cell proliferation and tumorigenicity

To unravel the roles of sORF1, we applied a gene editing strategy to induce sORF1 knockdown in SNU-16 cells (MsORF1) using the same guide RNA and strategy described above. The homologous recombination template (*pUC57-TUBA1B-MsORF1*) containing the *TUBA1B* exon 1 and intron 1 sequence in which the AUG codon of sORF1 was removed, and stop codons were introduced to disrupt sORF1, and the PAM sequence GGG was changed to GTG (Figure S5A). *pUC57-TUBA1B* with no sequence alteration except for the PAM was used as a negative control in unedited cells (WT). PCR plus restriction enzyme BsaAI digestion (Figure S5B), DNA sequencing (Figure S5C) and Western blot (Figure 3A) confirmed the successful gene editing of SNU-16 MsORF1 cells without disturbing α-tubulin. Notably, downregulation of CCND1 was observed in MsORF1 cells, indicating a potential deficiency in cell growth and proliferation (Figure 3A). Indeed, compared with the proliferation of wild-type SNU-16 cells, the proliferation of MsORF1 cells decreased by 25% (Figure S6), and MsORF1 cells exhibited a diminished ability to form colonies in a soft agar assay (Figure 3B). Moreover, MsORF1 cells were unable to form a stable monoclonal cell line. They coexisted with wild-type cells and were gradually overridden by wild-type cells as passages were carried on. To increase the portion of edited MsORF1 cells in the cell population, a pcDNA3.1-Neo plasmid was cotransfected into the cells during gene editing, and neomycin selection was applied. Thereafter, xenografts were generated in nude mice. WT and MsORF1 cells were inoculated subcutaneously into nude mice, and their tumour-forming ability was investigated. Tumours were palpable on day 10 and day 17 for the WT and MsORF1 groups, respectively (Figure 3C). Interestingly, in the MsORF1 group, two nude mice did not develop any tumours at all, while one mouse initially formed a tumour which subsequently completely regressed. The remaining three mice exhibited significantly smaller tumours compared to those formed by the parental WT cells. This indicates that the growth of tumours formed from MsORF1 cells was significantly lower than that of tumours formed from the WT cells (Figure 3D). Xenograft tumour sections showed positive sORF1 cells with no/low α-tubulin signal (Figure 3E), indicating the expression of TUBA1B-sORF1 *in vivo* and a similar small subgroup formed by the sORF1^+^/α-tubulin^low/-^ cells. To further examine whether the loss of proliferation and colony formation was due to the disruption of sORF1, reconstitution of sORF1 product was achieved using the rescue plasmid *pcDNA3.1-TUBRFP*, in which full-length *TUBA1B* cDNA was cloned with RFP tagged to the C-terminus of α-tubulin with no variation in the sORF1 sequence. Separately, *pcDNA3.1-TUBRFP-sORF1mFLAG* was constructed as a negative control, in which the sORF1 sequence was replaced with a 3XFLAG tag. Transfection of *pcDNA3.1-TUBRFP* partially rescued the CCND1 level and colony formation ability of MsORF1 cells (Figure 3F and 3G). These results suggest that TUBA1B-sORF1 contributes and is essential to cancer cell proliferation and tumour growth both *in vitro* and *in vivo*.

**Figure 3.**
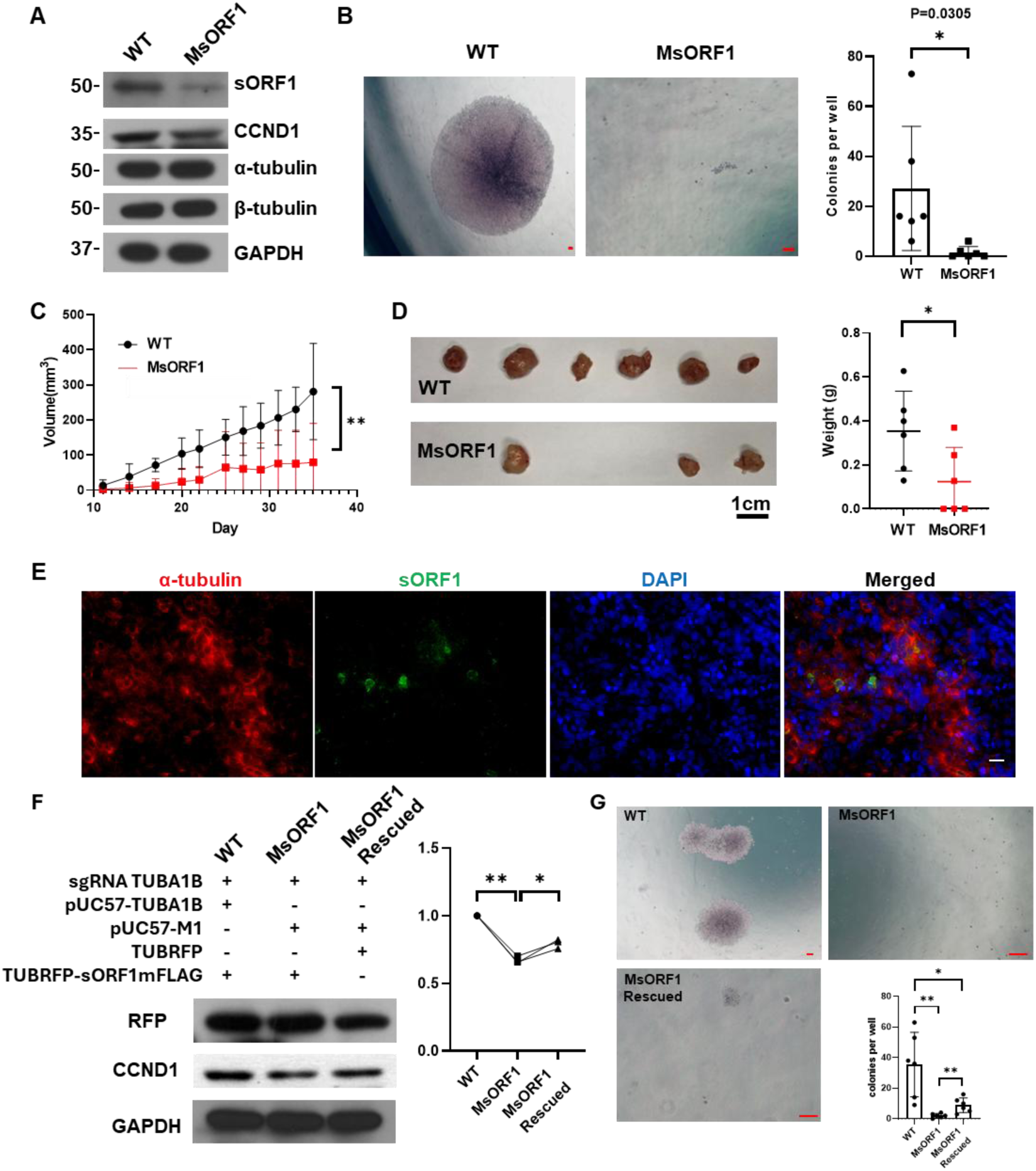
sORF1 is essential for cancer cell proliferation and tumour growth. (**A**) Cell lysates from wild-type (WT) and sORF1-knockdown SNU-16 (MsORF1) cells was applied with Western blot with the indicated antibodies. (**B**) WT and MsORF1 cells were seeded in soft agarose medium at 1x10^3^ cells/well in 24-well plates (6 wells each). The left panel shows representative images, and the right panel shows colony numbers on day 15. (**C-E**) SNU-16 WT and MsORF1 cells were xenografted into nude mice, followed by daily measurement of tumour size (**C**), collection of tumour tissue images taken on day 35 (**D**, left), statistical analysis of tumour weight (**D**, right) and immunofluorescence staining with the indicated antibodies (**E**). (**F, G**) Reconstitution of sORF1 in MsORF1 cells partially rescued CCND1 levels (**F**) and soft agar colony formation (**G**). sgRNA TUBA1B: TUBA1B guide RNA plasmid; pUC57-TUBA1B: wild-type DNA; pUC57-M1: mutant sORF1; TUBFRP: Full length of TUBA1B cDNA with RFP inserted in frame of α-tubulin. TUBFRP-sORF1mFLAG: TUBFRP plasmid with sORF1 was replaced with 3xFLAG. Scale bar: 100 μm; *: p<0.05; **: p<0.01.

### sORF1 participates in the transcriptional regulation of cell proliferation-related genes

To further elucidate the broader transcriptional alterations caused by the reduction in TUBA1B-sORF1, RNA sequencing was conducted on SNU-16 WT and MsORF1 cells. The transcriptional program of MsORF1 cells displayed broad reprogramming (Figure 4A). A total of 302 and 100 genes were downregulated and upregulated, respectively (Figure 4B), the majority of which were related to proliferation. The downregulated genes were proliferation-promoting genes, while the upregulated genes were proliferation-inhibitory genes. A full list of the changed genes is provided in Figure S7. The ontologies of the changed genes and their involvement in biological processes, cellular components and molecular functions (Figure 4C & Figure S8, Figure 4D & Figure S9) highlighted a broader reprogramming of the transcriptional processes in these cells as a result of the disruption of sORF1 coding, suggesting that sORF1 plays an important role in gene transcriptional regulation in cancer cells, especially for proliferation-related genes. Quantitative RT-PCR of a few representative genes confirmed the RNA-seq data (Figure S10).

**Figure 4.**
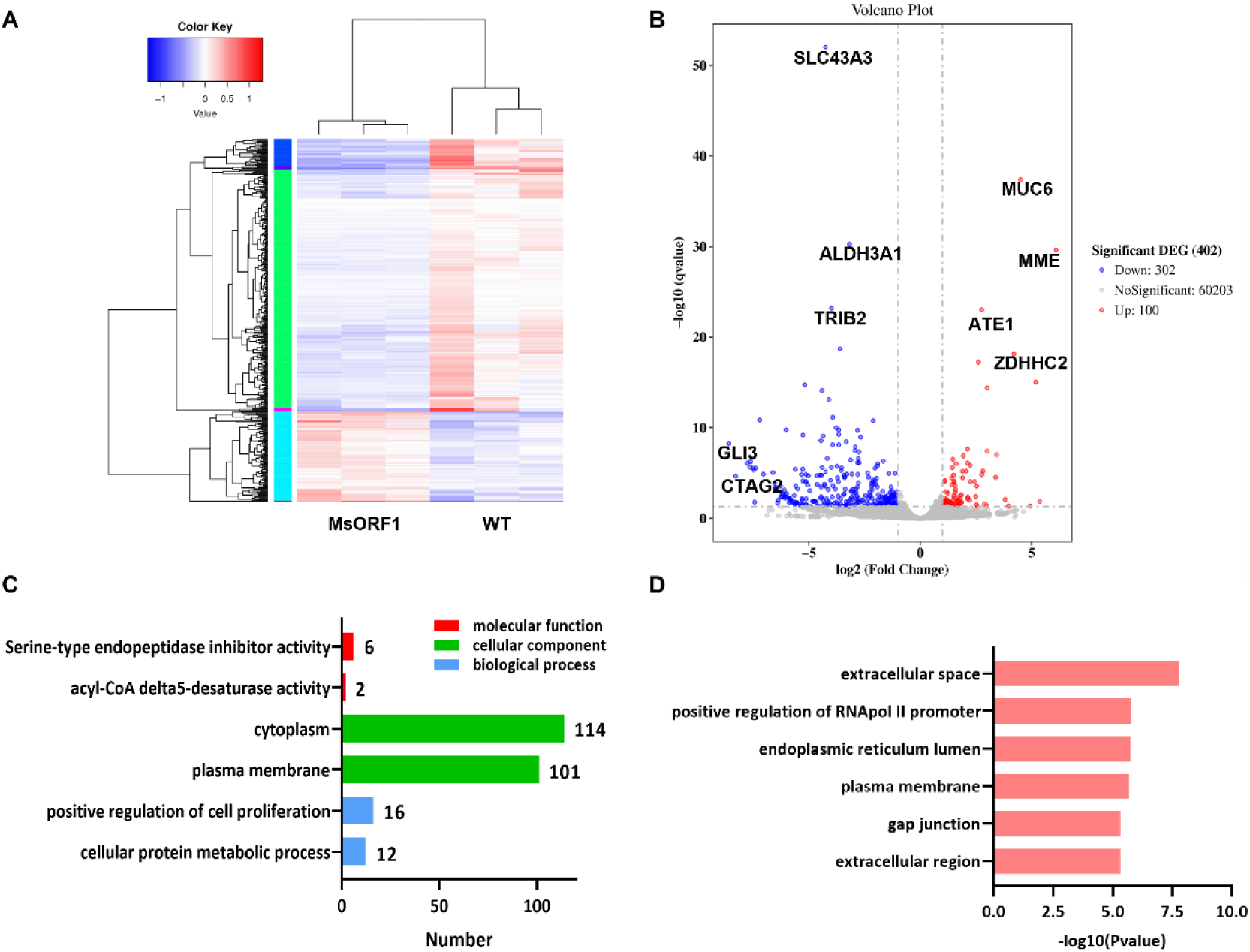
RNA sequencing analysis revealed 402 genes modulated by the disruption of TUBA1B-sORF1 in gene-edited SNU-16 MsORF1 cells. (**A, B**) Hierarchical clustering diagram (**A**) and volcano plot (**B**) of differentially expressed genes between SNU-16 WT and MsORF1 cells from three biological replicates. Downregulated and upregulated genes are indicated in blue and red, respectively. In the volcano plot, the x-axis represents the log2-fold change in gene expression, and the y-axis represents the negative log10 of the q-value. (**C**) Gene Ontology (GO) annotation and enrichment analysis of differentially expressed genes between SNU-16 WT and MsORF1 cells, where the y-axis represents the enriched GO term, and the x-axis represents the number of differentially expressed genes in that term. Different colours are used to indicate biological processes, cellular components, and molecular functions (the top two terms are shown). (**D**) Bar chart of the GO enrichment P value, with the y-axis representing the enriched GO term and the x-axis representing the negative log10 of the p value of that term (top six terms shown).

### sORF1 participates in protein nuclear translocation

Although disruption of sORF1 led to a significant decrease in cell proliferation and tumorigenesis, as well as the regulation of numerous crucial intracellular physiological processes, the specific mechanisms through which sORF1 affects cell behaviour remain unclear. As previously mentioned, Western blot analysis using anti-sORF1 antibodies detected multiple bands in cancer cells, and subsequent endogenous gene editing revealed various bands of FLAG-sORF1 products. These results suggest that the function of sORF1 may not solely rely on its own but rather on its involvement as a complex mediator in protein-protein interactions. Cellular fractionation combined with Western blot analysis revealed different sORF1 patterns between the cytosol and nucleus, with high-molecular-weight complexes dominating in the nucleus (Figure 5A), indicating a complicated role for sORF1 between the cytosol and nucleus.

**Figure 5.**
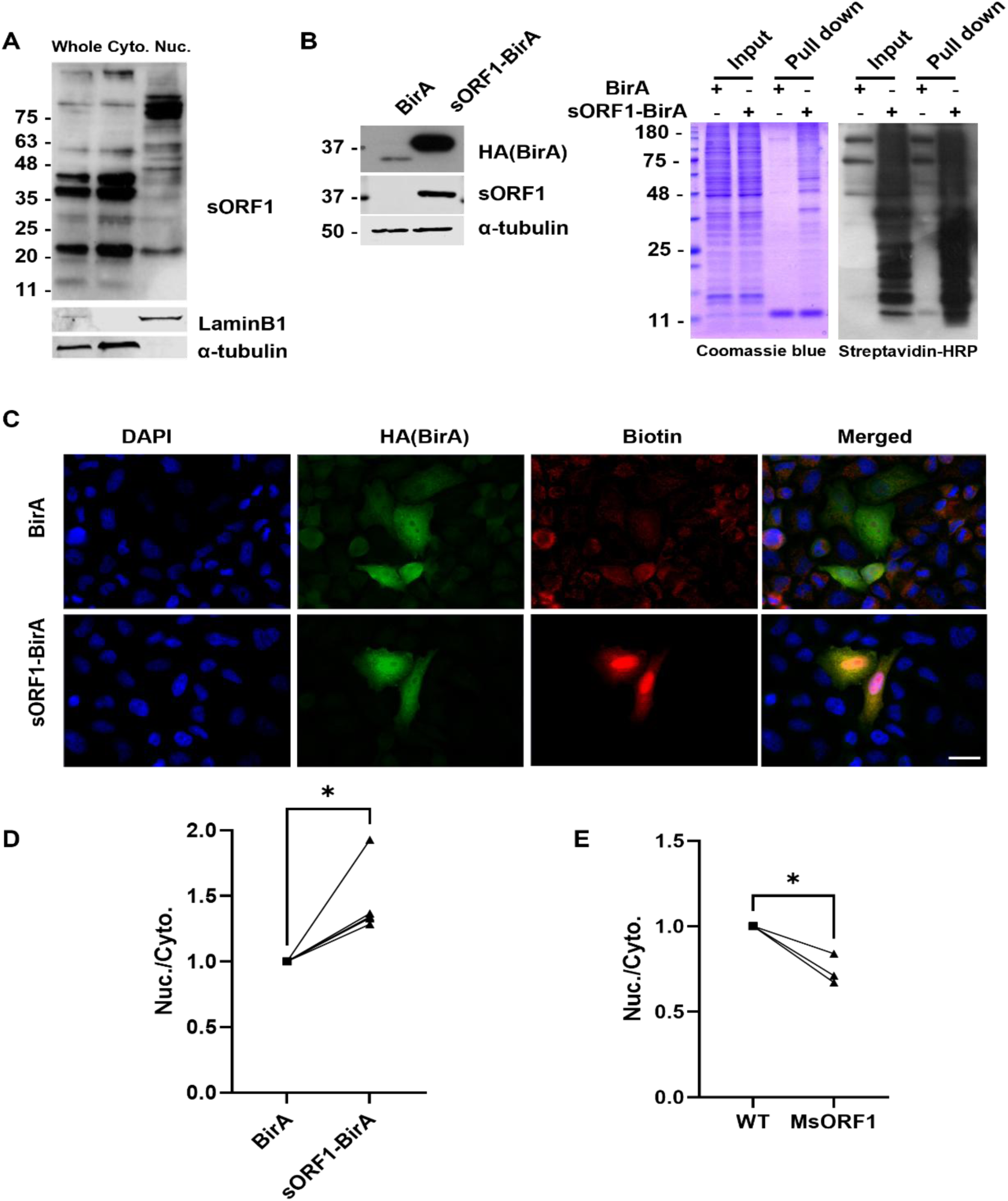
TUBA1B-sORF1 regulated nuclear protein transportation. (**A**) TUBA1B-sORF1 was differentially expressed in the cytosol and nucleus. Whole: whole-cell lysate; cyto: cytosolic fraction; nuc: nuclear fraction. LaminB1 and α-tubulin were included to indicate nuclear and cytosolic fractions, respectively. (**B-C**) HeLa cells were transfected with the pcDNA3.1-TUBA1B-sORF1-BirA(R118G)-HA vector, followed by WESTERN BLOT with the indicated antibodies (B, left); streptavidin pull-down with Coomassie blue staining and streptavidin-HRP detection (B, right); and IF staining of BirA (HA) and biotinylated proteins (fluorescein-labelled streptavidin) (C, left). (**D**) sORF1-BirA overexpression increased the nuclear protein concentration but decreased the cytosolic protein concentration. n=4 (**E**) Knockdown of TUBA1B-sORF1 (MsORF1 cells) led to downregulated nuclear proteins and upregulated cytosolic proteins concentration. n=3 *: p<0.05.

To investigate the protein-protein interactions involving sORF1, the system of proximity-dependent labelling of proteins, BioID, was applied to biotinylate the sORF1- associated protein in living cells where the sequence of sORF1 was inserted into *pc3.1-BirA(R118G)-HA* (Roux *et al*, 2012) to generate the vector *pc3.1-sORF1- BirA(R118G)-HA*, which encodes a fusion protein of sORF1-BirA(R118G)-HA (Figure 5B left). The biotin-labelled proteins covered a range from 10 kDa to 200 kDa within the *pc3.1-sORF1-BirA(R118G)-HA*-transfected HeLa cells, as revealed by Coomassie blue staining and Western blot with streptavidin-HRP (Figure 5B right), suggesting that sORF1 physically associates with a vast number of proteins. The biotinylated proteins were predominantly concentrated inside the nucleus, as revealed by IF staining with anti-HA and fluorescein-labelled streptavidin (Figure 5C). Intriguingly, quantification of the ratio of nuclear to cytosolic proteins revealed that overexpression of sORF1 via *pc3.1-sORF1-BirA(R118G)-HA* plasmid transfection increased the nuclear protein portion (Figure 5D), while sORF1 knockdown via the abovementioned gene editing decreased the nuclear protein portion (Figure 5E). Proteomic analysis of the streptavidin pull-down sample also revealed that sORF1-associated proteins were mainly located in the nucleus (Figure S11). These findings suggest that sORF1 regulates protein nuclear transport, thereby regulating gene transcription and influencing tumour cell proliferation and tumorigenicity.

### TUBA1B-sORF1 facilitates importin β-mediated β-catenin nuclear transportation

As described above, sORF1 disruption significantly inhibited the malignancy of cancer cells, accompanied by a decrease in the expression of CCND1, a downstream effector of the canonical Wnt/β-catenin pathway, at the mRNA and protein levels. Physical interactions were observed among sORF1, β-catenin (CTNNB1) and importin β (KPNB1), especially in the nucleus (Figure 6A, left). Importin β forms a transport cargo by binding to nuclear localization signal (NLS)-containing proteins, facilitating protein transportation into the nucleus. Despite lacking an NLS sequence, CTNNB1 can enter the nucleus by binding with KPNB1 (Fagotto *et al*, 1998). TUBA1B-sORF1 interacts with both CTNNB1 and KPNB1, suggesting that sORF1 may participate in KPNB1- mediated CTNNB1 nuclear translocation. Indeed, overexpression of sORF1 significantly increased CTNNB1 nuclear translocation but had no effect on KPNB1 (Figure 6A, right), while TUBA1B-sORF1 knockdown decreased CTNNB1 nuclear translocation (Figure 6B). Further experiments revealed that the absence of sORF1 attenuated the CTNNB1/KPNB1 interaction (Figure 6C), which could be restored with the same rescue strategy described above (Figure 6D). These results suggest that sORF1 may form a complex with CTNNB1 and KPNB1, facilitating CTNNB1 nuclear translocation, which in turn directs the transcription of genes involved in cell proliferation regulation.

**Figure 6.**
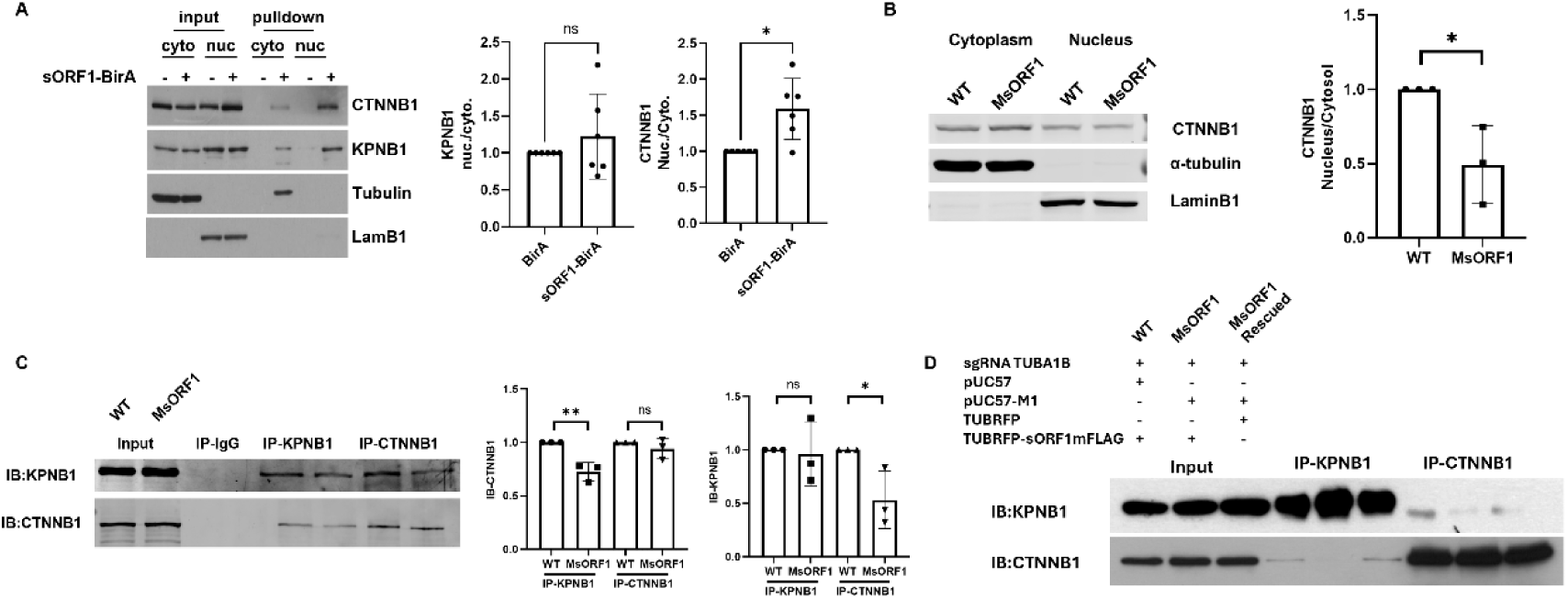
TUBA1B-sORF1 participated in importin β-mediated β-catenin nuclear translocation. (**A**) β-catenin and importin β were detected in sORF1-BirA streptavidin pull-down samples (A, left), where β-catenin nuclear translocation significantly increased while importin β did not significantly change (A, right). n=6 (**B**) Nuclear translocation of β-catenin was decreased in TUBA1B-sORF1-knockdown HeLa cells (MsORF1). n=3 (**C-D**) The interaction between β-catenin and importin β was disrupted in MsORF1 cells (**C**) and was rescued by reconstitution with TUBA1B-sORF1 (D). ns: not significant. n=3. *: *p*<0.05, **: *p*<0.01.

### The sORF1 protein is upregulated in human gastric cancer tissues and serum

As described above, TUBA1B-sORF1 was essential for tumour growth in the xenograft model. Further immunohistochemistry (IHC) staining was performed on formalin-fixed paraffin-embedded sections from 10 groups of gastric carcinoma patients, and both the tumour and paratumour tissues were analysed. A small proportion of sORF1- positive cells were detected in both tumour and paratumour tissues (Figure 7A). Semiquantitative analysis revealed a greater percentage of sORF1-positive cells in tumour tissues than in paratumour tissues, but the difference was not significant, possibly due to the small number of samples (Figure 7B-C). However, when tumour grade was evaluated (Figure 7D), a significant increase in sORF1-positive cells was detected in the paratumour area in the undifferentiated-grade tumour group compared to the differentiated-grade group (Figure 7D). Furthermore, an enzyme-linked immunosorbent assay (ELISA) revealed that there was a significantly higher level of serum sORF1 in cancer patients than in healthy individuals (Figure 7E). As expected, a subpopulation of sORF1^+^/α-tubulin^low/-^ cells was also detected (Figure 7F, upper). Further experiments revealed the coexistence of sORF1 and OCT4, a stem cell marker, in gastric carcinoma tissues (Figure 7F, lower). These observations suggest a translation switch from α-tubulin to sORF1 similar to that observed in the *in vitro* study and indicate the potential of these sORF1^+^/α-tubulin^low/-^ cells as putative cancer stem cells. Overall, these results highlight the potential of sORF1 as a novel therapeutic target and prognostic biomarker for cancer treatment.

**Figure 7.**
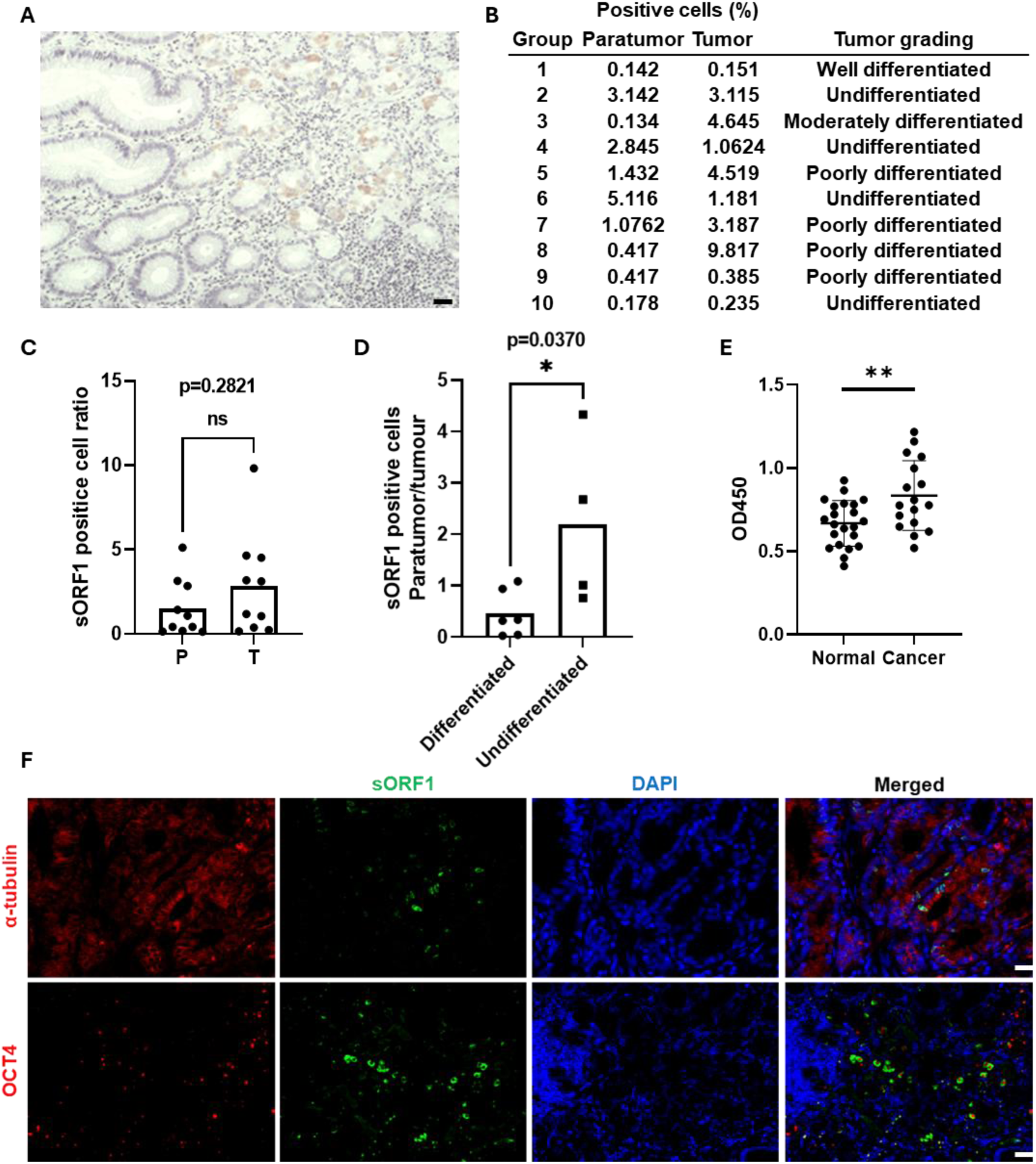
The sORF1 protein contributes to tumorigenesis. (A) Representative images of immunohistochemical staining of sORF1 in gastric carcinoma tissue sections. (B) Comparison of sORF1-positive cells between paratumour (P) and tumour (T) tissues. (C) The percentage of sORF1-positive cells in the ten groups of gastric carcinoma tumour and paratumour sections according to tumour grade. (D) Comparison of sORF1-positive cells between differentiated and undifferentiated tumours. *: p<0.05. (E) Enzyme-linked immunosorbent assays revealed a higher level of serum sORF1 in cancer patients (n=17) than in healthy individuals (n=21). **: p<0.01. (F) Representative images of IF staining with the indicated antibodies. DAPI was used to counterstain the nuclei. Scale bar: 100 μm.

## Discussion

In this study, we demonstrated for the first time that an sORF can be alternatively translated from the *TUBA1B* gene via both canonical methionine and noncanonical leucine initiation in cancer cells, the product of which, TUBA1B-SORF1, functions independently of its mORF product, the α-tubulin 1b protein. Remarkably, disruption of sORF1 translation significantly reduced cancer cell proliferation *in vitro* and tumorigenesis *in vivo*. Subsequent investigations revealed that sORF1 participates in protein nuclear translocation regulation, an example of which is importin-β-mediated β-catenin nuclear translocation. Consequently, the transcription of genes involved in cell proliferation is modulated.

Tubulin is an evolutionarily conserved superfamily of structural proteins comprising six families (α-, β-, γ-, δ-, ε- and ζ-tubulins) (Nsamba & Gupta, 2022). Tubulin exerts diverse functions in multiple cellular processes, including structural support, intracellular cell transportation and DNA segregation. Mammalian cells possess α-, β- and γ-tubulin. The former two proteins form heterodimers, the block unit of microtubules with the α-subunit at the minus end and the β-subunit at the plus end. γ- tubulin is responsible for the nucleation and polar orientation of microtubules. β- subunit-bound GTP undergoes the GTP/GDP cycle, an essential process for the dynamic instability of microtubules. The β-subunit is the target of all MTA drugs. α- subunit-bound GTP does not undergo hydrolysis and therefore stabilizes the β-subunit. More variation at the mRNA and protein levels has been reported for α-tubulin than for β-tubulin in cancer cells (Hu, 2021; Xiong, 2019; Qin, 2021; Linge, 2012; Wang *et al*, 2020). In this study, similar phenomena were observed in gastric and colorectal cancer tissues. There are eight isoforms of α-tubulin, which are derived from different alleles. Different alleles may be expressed in different tissues or cells; for example, *TUBA1A* is related to neurogenesis (Aiken *et al*, 2019). The discrepancy between the mRNA and protein levels of the *TUBA1A* and *TUBA1B* genes have been reported (Wang *et al*, 2020; Xu, 2020). The variation in the α-tubulin protein can theoretically be derived from either translational regulation or posttranslational stability. Accumulating evidence indicates that the stability of α-tubulin can be regulated by PTMs via acetylation, phosphorylation, ubiquitination, etc (Gadau, 2019; Roth, 2022; Xu *et al*, 2010). The contribution of translational regulation to the variation in the α-tubulin protein level remains underestimated.

Great progress in ribosomal profiling-derived translatomics and proteomics has led to the identification of a diverse range of sORFs in mammalian cells (Leong, 2022). The spatiotemporal translation of sORFs may be involved in different physiological and pathophysiological processes, including cancer. In this study, the proteomics assays detected peptide fragments from TUBA1B-sORF1 but not from other alleles. An experiment involving the insertion of the mCherry sequence confirmed the translation of sORF1 from *TUBA1B* but not from *TUBA1A*, though the AUG codon of *TUBA1A- SORF1* surpasses the AUG codon of the mORF for α-tubulin 1A. Although the GWIPS-viz Genome Browser analysis (Kiniry *et al*, 2018) for *TUBA1B* did not suggest the translation of sORF1 from the canonical methionine codon, the strategy with the insertion of mCherry and the Met^AUG^/Ile^AUC^ mutation indeed confirmed the translation of *TUBA1B-sORF1* from the methionine codon, which reminds us to take caution when interpreting data from ribosomal profiling. The majority of the cancer cell population was positive for both TUBA1B-sORF1 and α-tubulin according to IF staining, indicating the constitutive translation of sORF1 and the dual-cistronic nature of the *TUBA1B* gene. The sORF1 sequence displays high homology among different alleles of TUBA1 genes, but TUBA1B-sORF1 possesses a unique C-terminus compared to that of sORF1 in other alleles. Whether this unique C-terminal sequence is involved in its own translation requires further detailed investigation. Although the translation of sORF1 to other alleles was not supported in cancer cells in this study, its translation to other cell types or tissues cannot be excluded.

Noncanonical translation or alternative translation widely exists in prokaryotic genomes to increase genome capacity (Cao & Slavoff, 2020). Recent progress in proteogenomic and ribosomal profiling has revealed that non-AUG codon-initiated translation, especially in sORFs, widely occurs in eukaryotic genomes, including mammalian genomes (Andreev, 2022; Green *et al*, 2022; Kearse & Wilusz, 2017). An increasing number of genes that can produce not only classical protein or main open reading frame (mORF) products but also overlapping alternative open reading frame (ORF) or short open reading frame (sORF) products with completely different amino acid sequences have been discovered. These genes have been named dual-coding genes due to their ability to encode 2 distinct proteins from the same sequence (Cao, 2021 ;Khan, 2020). Previous studies of dual-coding genes reported that the alternative products from POLG and RPL36 also used CTG and GTG instead of AUG as start codons for translation (Khan, 2020). Translation initiation from non-AUG codons may involve canonical Kozak sequences (Schwab *et al*, 2004), internal ribosome entry sites (Touriol, 2003) or chemical modifications of codons (Fujita, 2022). This alternative translation initiation appears to prefer non-AUG codon-guided initiation. In this study, the proteomics data and the strategies using 3xFLAG tag insertion and mutagenesis demonstrated that the translation of sORF1 can be initiated partly from three upstream non-AUG codons, one of which, the Leu codon (94) of sORF1, colocalizes with the Met codon (96) of the mORF for α-tubulin. There may be codon selection between Leu for sORF1 and Met for the mORF when the ribosome binds to *TUBA1B* mRNA and initiates translation. This selection may account for the subpopulation of sORF1^+^/α- tubulin^low/-^. The translation shift between sORF1 and mORF can provide another option for the variation of the α-tubulin protein in addition to its stability. Initiation from the non-AUG codons produces sORF1 with an extended N-terminus, which can be post translationally modified via serine residues or the cysteine residue from Leu (94), increasing the molecular weight compared with what was expected. Further detailed investigation is required to verify the other two non-AUG codons, the PTM sites and types, and the functional comparison among these isoforms and the canonical sORF1 product.

The canonical translation of sORF1 produces a microprotein of 50 amino acids with an apparent molecular weight of 6 kDa. The gene-edited FLAG insertion (<11 kDa band in Figure 2C), sORF1-mCherry fusion protein (Figure 1C and Figure 2F) and sORF1- BirA (HA) fusion protein (Figure 5B) displayed the expected molecular weight. However, sORF1 antibody detection of endogenous sORF1 and FLAG antibody detection of FLAG-tagged sORF1 revealed multiple bands with higher molecular weights. The ablation of the bands by peptide blocking indicates the antigen specificity of the antibody. These bands can be either derived from the PTM of the sORF1 product, as observed in the non-AUG codon-initiated product, or most likely from sORF1 serving as a PTM donor itself, such as through ubiquitination (Pickart & Eddins, 2004) and ubiquitin-like modification (SUMOylating) (Johnson, 2004), in which the C-terminus is important for this sORF1-mediated modification. Fusion to mCherry or BirA(HA) blocks the access of the C-terminus to target substrates, increasing the expected size of the fusion proteins. FLAG-tagged protein has a free C-terminus, giving rise to multiple bands. Supporting evidence also came from an additional experiment (Figure S12). N-terminally biotinylated sORF1 (b50A in Figure S12) bound more proteins than C-terminally biotinylated sORF1 (56Ab in Figure S12, the upstream 6 aa included), as revealed by Western blot analysis with anti-SORF1 and streptavidin-HRP (a horseradish radish peroxidase). SUMOylating mainly regulates substrate cellular location (Johnson, 2004). In this study, the sORF1 protein level was positively related to protein nuclear translocation. Thus, sORF1-mediated modification may also affect the cellular location of the substrate. However, further detailed investigations on this topic are needed.

α-tubulin plays a critical role in cellular transportation and cell mitosis. In this study, we found that sORF1 plays a similarly important role in these processes. The sORF1 protein level is positively associated with protein nuclear translocation. Overexpression of sORF1 increased protein nuclear translocation, while disruption of sORF1 decreased protein nuclear translocation, which could be rescued by reconstitution of wild type sORF1. Furthermore, disruption of sORF1 abolished cancer cell proliferation, as revealed by soft agar colony formation, which could also be partly rescued by reconstitution of wild-type sORF1. The tumorigenesis of MsORF1-edited cells must be derived from coexisting wild-type cells. Therefore, the translation of sORF1 and mORF coexists in most cells. Although no significant overlap of signals from sORF1 and α- tubulin was detected in double IF staining, the association of sORF1 and α-tubulin was indeed detected in BioID (Roux *et al*, 2012) studies, suggesting that there may be some links between these two proteins from the same *TUBA1B* mRNA similar to that observed in Ataxin-1 (Bergeron, 2013). As sORF1 exists in both the cytosol and nucleus, while α-tubulin exists exclusively in the cytosol, sORF1 and α-tubulin can work independently, similar to MUC1-ARF versus MUC1-TM (Chalick, 2016). There is a correlation between sORF1-mediated protein nuclear translocation and cell proliferation, and the effective mediators are proliferation-related genes. An example supporting this notion is the interaction of sORF1 with the Wnt signalling pathway. sORF1 forms a complex with importin-β and β-catenin, facilitating β-catenin nuclear translocation and thereby increasing cyclin D1 expression. There was a positive relationship among the sORF1 protein level, β-catenin nuclear translocation and cyclin D1 expression. Further investigation is required to verify whether upstream Wnt ligand binding is needed for sORF1 translation or whether sORF1 is translated intrinsically and activates Wnt signalling in a Wnt ligand-independent manner.

In clinical research, the expression pattern of sORF1 in gastric cancer tissue differs from that observed in cell lines, where sORF1-positive cells are present but in limited quantities. Tumour cells exhibit heterogeneity, and recent single-cell-based studies have identified the potential existence of cancer stem cells/cancer-initiating cells and metastasis-initiating cells within the bulk tumour population (Massague & Ganesh, 2021; Bonnet & Dick, 1997; Al-Hajj *et al*, 2003; Singh, 2003; Shimokawa, 2017). Cancer stem cells are believed to be “seeds” for tumour growth and metastasis. Distinguishing these distinct cell populations can aid in developing therapies specific to subgroups of cancer cells (Takebe, 2015; Laszlo *et al*, 2014). In this study, we demonstrated the essentiality of sORF1 translation for tumour growth via *in vitro* soft agar colony formation assays and an *in vivo* xenograft model. In cancer tissues, sORF1 is coexpressed with the cancer stem cell marker OCT4 in the same cells. These observations suggest that there may be a direct link between sORF1 translation and cancer stem cell regulation. Further detailed investigations on this link could unlock new avenues for the development of targeted therapies and improve treatment strategies for cancer patients. Additionally, the presence of elevated levels of sORF1 in serum samples from tumour patients suggests that sORF1 may serve as an early diagnostic biomarker for some types of cancer.

In summary, this study uncovers novel regulatory mechanisms involved in *TUBA1B* gene mRNA translation. *TUBA1B* is a dual-cistronic gene encoding α-tubulin from the mORF and a microprotein from the sORF1. Both ORFs can be translated in the same cell, and there is a translation switch between them, leading to variations in the α- tubulin protein level. sORF1 can be translated in a methionine-initiated canonical manner or a leucine-initiated noncanonical manner. sORF1 serves as a chaperone or a PTM modifier to facilitate protein nuclear transportation, subsequently regulating proliferation-related gene transcription and leading to cancer cell proliferation and tumour growth. There may be a link between sORF1 translation and cancer stem cell regulation. This study provides new insights into the molecular mechanism underlying cancer development and highlights new potential targets for therapeutic intervention in cancer treatment and biomarker for cancer diagnosis.

## Materials and methods

### Cell Culture

The HEK-293 (CRL-1573), HeLa (CCL-2) and SNU-16 (CRL-5974) cell lines were purchased from ATCC and maintained in DMEM (HEK-293 and HeLa) or RPMI 1640 (SNU-16) supplemented with 5% fetal bovine serum (FBS, Gibco) and penicillin/streptomycin in a humidified 37°C incubator with a 5% CO_2_ supply.

### Xenograft Models

All animal experiments were performed at Shanghai CroChinese Biotechnology Co., Ltd. (Shanghai, China) according to protocols approved by the Institutional Committee for Use and Care of Laboratory Animals under the Chinese guidance GB/T35892-2018. Eighteen 4- to 5-week-old female BALB/C nude mice were purchased from Hangzhou Ziyuan Experimental Animal Technology Co., Ltd. (Hangzhou, China). After acclimation for one week, 12 nude mice were randomly divided into two groups, and 5x10^6^ cells in 100 µl of PBS containing SNU-16 or SNU-16-MsORF1 cells were injected subcutaneously into the dorsal neck of each mouse. The maximum and minimum tumour sizes were measured with callipers, and the body weights of the mice were measured every other day. On day 35, the mice were sacrificed humanely via dislocation of the neck. All the tumour tissues were isolated, followed by image collection and FFPE treatment.

### LC-MS/MS peptide identification/Proteomic analysis

Cell lysates, immunoprecipitates or biotin-streptavidin bead-mediated pull-down samples were separated by SDS-PAGE, fixed and stained with Coomassie blue solution (Bio-Rad). The bands were cut and subjected to in-gel trypsin digestion and tandem liquid chromatography-mass spectrometry. The data were viewed and analysed by Scaffold (v5.1.2).

### Cellular fractionation

The cytosolic and nuclear fractions were isolated using a modified standard procedure described elsewhere. Briefly, the cells were resuspended in 100 μl/10^6^ hypotension (10 mM Tris-HCl, pH 7.5, 10 mM KCl plus protease inhibitors) buffer and incubated on ice for 15 minutes with agitation every 5 minutes. At a final concentration of 0.625%, NP-40 was added, and the mixture was vortexed at the highest speed for 10 seconds to disrupt the swollen plasma membrane. The cell suspension was centrifuged at the highest speed for 10 seconds to pellet the nuclei. The supernatant was collected as the cytosolic fraction. The nuclei were washed once with PBS and then resuspended in 100 μl/1x10^6^ hypotension buffer containing 0.625% NP-40, followed by sonication (power set at 1) for several seconds and subsequent incubation on ice for 30 minutes. The supernatant was collected as the nuclear fraction after centrifugation at the highest speed for 5 minutes. All centrifugation steps were performed at 4°C. The cytosolic and nuclear fractions were subjected to Western blot or immunoprecipitation after protein concentration quantification with Bito-Rad reagents.

### Western blot, immunoprecipitation, and immunofluorescence staining

Western blot, immunoprecipitation and immunofluorescence staining were performed according to standard procedures described elsewhere. For Western blot and IP, the cells were lysed in IP-A buffer (10 mM Tris-HCl, pH 7.5, 120 mM NaCl, 1 mM EDTA, 1% Triton X-100 plus protease inhibitors). The protein concentration was measured using Bradford reagents (Bio-Rad). Twenty-five micrograms of protein was used for Western blot. For IP, 1 mg of cell lysate was incubated with 2 µg of antibody or normal IgG, 2 volumes of IP-B buffer (IP-A without Triton X-100) and 5 µl of protein G magnetic beads on a rotator at 4°C overnight to pull down the target and its associated proteins, followed by Western blot or proteomics analysis. For IF, cells were directly fixed with 4% paraformaldehyde in PBS and then permeabilized. FFPE tissue samples were subjected to dewaxing and antigen retrieval, followed by permeabilization and subsequent standard steps. Fluorescence was observed under a ZEISS Axio Vert. A1 and LAS AF confocal microscopy. For the peptide blocking assay in Western blot and immunofluorescence staining, the anti-sORF1 antibody was preincubated with antigen peptide (1:10) in blocking buffer overnight before being applied to the samples. (Antibodies list - Supplementary Table S1)

### Immunohistochemistry staining and semiquantification

FFPE sections were placed in a heat incubator at 60°C overnight, subsequently dewaxed using xylene, and rehydrated through a series of ethanol concentrations (100%, 90%, 80%, and 70%) for 5 minutes each. Next, 0.1 M sodium citrate (pH 6.0) was used for antigen retrieval. A Solarbio IHC SP kit (SP0041) was used for subsequent procedures according to the manufacturer’s guidelines. The primary antibody, anti-sORF1, was diluted to a concentration of 1 µg/ml, and the blocked anti- sORF1 was preincubated with an immunogen peptide at 10 µg/ml at 4°C overnight. sORF1-positive cells were enumerated using Halo v3.0.311.314 - Multiplex IHC v2.2.0 software (Indica Labs).

### Enzyme-linked immunosorbent assay

Single wells of a 96-well microplate were coated with a mixture of 2 µl of human serum and 98 µl of CBB coating buffer and incubated overnight at 4°C. Subsequently, the wells were washed four times with PBST, followed by blocking with 5% skim milk/PBST solution. The subsequent procedures were carried out using an ELISA kit (Solarbio SEKF105) according to the manufacturer’s instructions. The anti-sORF1 antibody was diluted to a concentration of 1 µg/ml. Each sample was analysed in triplicate to ensure technical reproducibility.

### Flow Cytometry

The SNU-16 cells were collected by five-minute centrifugation at 300 g, fixed with 4% PFA for 15 min at room temperature, and then permeabilized with 0.1% Triton X- 100/PBS for another 15 min at room temperature. Next, the cells were blocked with 10% normal goat serum/PBS for 1 hour at room temperature, followed by incubation with primary antibodies overnight at 4°C (antibodies list - Supplementary Table S1). After primary incubation, the cells were washed with ice-cold PBS three times then incubated with FITC- or Cy3-conjugated secondary antibodies for 1 hour at room temperature, followed by washing with PBS. The fluorescently labelled cells were filtered through a 70 μm cell strainer and analysed with a Beckman DxFLEX Flow Cytometer. (Gating strategy – Supplementary S4 and S5).

### Soft Agarose Colony Formation Assay

0.7% and 1% (w/v) agarose solutions were autoclaved and equilibrated to 40°C. Then, 1% agarose solution was mixed with an equal volume of 2× RPMI-1640 media supplemented with 2× penicillin/streptomycin and 20% FBS. One milliliter of the mixed media was added to a 24-well plate and allowed to solidify for 10 min to form the bottom layer. Then, 0.7% agarose solution was mixed with 2X media, and SNU-16 cells were added to a final concentration of 1,000 cells/ml. One milliliter of the cell/agarose/media mixture was plated on the bottom layer. The plate remained at room temperature for 10 minutes to achieve complete gelation. Then, 1X RPM1-1640 supplemented with 1X penicillin/streptomycin and 10% FBS was added to each well. The cells were cultured for 3 weeks while the liquid media was refreshed twice a week.

### Plasmid construction and transfection

The construction and verification services for all the plasmids used in this study were provided by Beijing Biomed Gene Technology (Beijing, China), including gene synthesis, FLAG/mCherry (red fluorescence protein, RFP)/GFP (green fluorescence protein) tag insertion, site mutagenesis and DNA sequencing. Plasmid DNAs were amplified and purified from JM109 competent cells using Qiagen miniprep kits. The plasmid DNAs were transfected into HEK293, HeLa or SNU-16 cells using Xfect polyers according to the manufacturer’s protocol. The fluorescent proteins were observed under a Keyence BZ-X710 microscope.

### CRISPR/Cas9-homologous recombination-based gene editing in SNU-16 cells

The guide RNA sequence GTGACCACCTAGTCTAAAGT was utilized to generate the pX330-TUBA1B-sgRNA plasmid. The TUBA1B gene sequence (NM_006082.3) with specific alterations, including the insertion of a FLAG tag upstream of the AUG in the sORF for FLAG labeling or nucleotide modifications to eliminate the AUG start codon and introduce early stop codons to disrupt sORF1, was used to create the recombinant plasmids. These constructs were designated as pUC57-FsORF1 (for FLAG insertion) and pUC57-MsORF1 (for sORF1 disruption), respectively. The PAM sequence of GGG was changed to GTG in pUC57-FsORF1 and pUC57-MsORF1. These plasmids were created and verified by Beijing Biomed Gene Technology (Beijing, China). SNU-16 cells were transfected with pX330-TUBA1B-sgRNA, pCI-Neo and pUC57-FsORF1 or pUC57-MsORF1 and selected by geneticin.

### DNA extraction, PCR and DNA sequencing

The gene edited transfected cells were resuspended in lysis buffer (50 mM Tris-HCl pH 8.0, 100 mM EDTA, 100 mM NaCl, 1% SDS, and 1 mg/mL proteinase K) and mixed with one-third volume of 3 M sodium acetate (pH 6.5), followed by centrifugation to remove precipitates. The supernatant was recovered and mixed with the same volume of isopropanol, followed by centrifugation to precipitate the DNA pellet. The pellet was washed once with 70% ethanol and then resuspended in DNase-free water. The concentration was checked by Nanodrop (Thermo Fisher) and adjusted to 20 ng/µl if necessary. Twenty nanograms of DNA was subjected to PCR with Premix Taq plus dye (Takara, RR901A) according to the manufacturer’s instructions.

The following primer set was used to generate a specific 499 bp product by PCR from SNU-16-FsORF1 cells: FsORF1_F: gattctgcagtacagatgtcc versus FsORF1_R: cgtcgtcgtccttgtaatctg.

The following primer set was used to generate a specific 665 bp product by PCR from both SNU-16 WT and SNU-16-MsORF1; MsORF1_F: agactggtagtaggccaatggctg versus MsORF1_R: gagttgctcagggtggaagagctg. The PCR products from MsORF1 cells will be digested by the restriction enzyme BsaAI (NEB, R0531S).

The following primer set was used to amplify the product covering the mutation sequence, and then the product was subjected to Sanger sequencing. MsORF1_F2: agactggtagtaggccaatggctg versus MsORF1_R2: gagttgctcagggtggaagagctg.

### Biotin-labelled peptide/streptavidin pulldown

HeLa cells were collected and washed with PBS and then lysed in PBS containing 1% Triton X-100 and protease inhibitors, followed by 5 seconds of sonication at a power of 1.0. The supernatant was collected, and the total protein concentration was measured and then adjusted to 1 mg/ml with PBS. One milliliter of protein supernatant was mixed with 20 μl of streptavidin magnetic beads and incubated on a rotator at 4°C for 1 hour to preclear endogenous biotin-labelled proteins. The supernatant was recovered and incubated with 2.5 μg of biotin-labelled sORF1 proteins in a water bath at 37°C for 1 hour. The following steps were also used to isolate sORF1-associated proteins from sORF1-BirA-transfected cells. Twenty microliters of streptavidin magnetic beads were then added and incubated on a rotator at room temperature for 1 hour. The beads were trapped with magnetic bar and washed with PBS containing 1% Triton X-100, resuspended in 100 μl of 1X SDS loading solution containing 0.2 M DTT and incubated at 95°C for 10 minutes. The samples were subjected to Western blot or proteomics analysis.

### RNA sequencing and RT-qPCR

For RNA sequencing, SNU-16 and SNU-16-MsORF1 cells were lysed in TRIzol reagent and sent to Azenta (Suzhou, China) for subsequent sequencing. Total RNA was isolated with Qiagen miniprep kits and reverse transcribed using ImProm-II reverse transcriptase (Promega, A3802) according to the manufacturer’s protocol. The plates were diluted 1/10 and subjected to qPCR using qPCRBIO SyGreen Blue Mix (Hi-ROX) (PCR Biosystems, PB20.16-01) on an Eppendorf mastercycler. The following primer sets were used to verify the significantly differentially expressed genes from the RNA sequencing results.

NUP210_F: gctaccgctggttgtcca versus NUP210_R: tggatgaggtccacaatggc; H19_F: caaagc ctccacgactctgt versus H19_R: tggggcgtaatggaatgctt; MUC6_F: cccctgccttctcctctcag versus MUC6_R: ctgcagaccccgggtga; ATE1_F: tactgcaagaacgagtcggg versus ATE1_R: ccatgggctc atcctcacaa; LGI2_F: tgacgtgaccagctttgact versus LGI2_R: cccacgatggactgacctgt; TUBA1B_F: tcgcctcctaatccctagcc versus TUBA1B_R: gccagtgcgaacttcatcaat; GAPDH_F: aatgggca gccgttaggaaa versus GAPDH_R: gcccaatacgaccaaatcagag.

### Gastric Carcinoma FFPE Sections

The human gastric and colorectal carcinoma tissues and formalin-fixed paraffin-embedded (FFPE) sections used in this study were obtained from the Chinese Natural Population Biobank of Ningbo No.2 Hospital, Ningbo, China. The study has passed and followed the ethical review from the Human Research Ethics Committee of Ningbo No.2 Hospital (YJ-NBEY-KY-2021-125-01).

## QUANTIFICATION AND STATISTICAL ANALYSIS

The data are presented as the means ± SDs as indicated, and statistical significance was determined using two-tailed Student’s t tests. Representative images of three biological replicates are shown for the Western blot results. Densitometric analysis of the bands was performed with ImageJ, and the results are presented as the mean ± SD. Statistical analysis was performed using two-tailed Student’s t tests, and a significance level of p < 0.05 was considered to indicate statistical significance.

## AUTHOR CONTRIBUTIONS

L.Z. and T.C. conceived and designed the project. X.B. and Y.T. performed the experiments, analysed the data and wrote the manuscript. Y.Z., Y.Z., L.Y., and S. Z. performed the experiments; X.H., E.S. and M.G. designed the experiments; H.L., A.Z., A.M., and J.N.A. performed the critical review of the manuscript.

