## Supplementary figures for "Human TUBA1B short open reading frame product regulates cancer cell growth via importin β"

| | | |
|---|---|---|
| Homo Sapiens | **TUBA1A** | MPAGSSTAWNTASSPMARCQVTRPLGEEMIPSTPSSVRRGLASMCPGQCL |
| | **TUBA1B** | MPAGSSTAWNTASSPMARCQVTRPLGEEMTPSTPSSVRRALASTCPGLCL |
| | **TUBA1C** | MPAGSSTAWNTASSPMARCQVTRPLGEEMIPSTPSSVRRGLASMCPGQCL |
| | **TUBA3C** | MPAGNCTAWNMEFSPMVRCQVIKPLVVGTTPSTRSSVRLELASTCPEQCLW<br>TWSPLWSMKCAQEPIGSSSTQSS |
| | **TUBA3D** | MPAGNCTALNMEFSPMAKCQVIKPLVAGTTPSTRSSVRLELASTCPEQCLW<br>TWSPLWSMKCAQGPTGSSSTRSS |
| | **TUBA3E** | MPAGNCTALNMEFSPMVKCQVIKPLVAGTTPSTRSSVRLELASTCPEQCLW<br>TWSPLWSMKCAQGPTGSSSTQSS |
| | **TUBA4A** | MPAGSSIAWNMGFSLMGRCPVTRPLVEGTTPSPPSSVKLVLENTYPGQFL<br>WIWSLRSLMRSEMAHTDSSSTQSSSSLGKRMLPTTMPVVTIPLARRSLTQ<br>CWIGSASCLTSAQDFRASWCSTALVGALALASPHS |
| | **TUBA8** | MPAGSSSAWNTASRQTALLMLKLARSTMMTPSPPFSARLAMGSMCPGP<br>S |
| Mus Musculus | **TUBA1A** | MLARLVSRSAMPAGSSTAWNMASSLMARCQVTRPLGEEMTPSTPSSVRQE<br>LASMCPGQCS |
| | **TUBA1B** | MPAGSSTAWNMASSLMARCQVTRPLGEEMTPSTPSSVRQELASMCPGQCS |
| | **TUBA1C** | MPAGSSTAWNMASSLMARCQVTRPLGEEMTPSTPSSVRQELASMCPGQCS |
| | **TUBA3A** | MPAGNCTALNMAFSLTVRCQATKPLAAGTTHSTHSSVRLEPASTCPGQCLW<br>TWSPLWWMRCAREPTGSFFTQSS |
| | **TUBA3B** | MPAGSCTALNMVFSLTVRCQATKPLAAGTTHSTHSSVRLEPASTCPGQCL<br>WTWSPLWSMKYAPEPTGSFSTQSSSSLGRKMQPTIMPEVIIPLARRLLTW<br>SWTGSENWPICARDCRASSSSTALEVAQGLGLHRC |
| | **TUBA4A** | MWGRQVSRWAMPAGSSTVWNMGFSLMGRCPAIRPLVEGTTPSPPSSVKLE<br>LENMCLGQSLWTWSLL |
| | **TUBA8** | MWVKPEFRLAMPAGSSSAWSTASRRMEPLALRPARSTMTTPLPPSSVRLA<br>TGNTCPGLSWWTWSLP |

**Figure S1. A schematic illustration of the sORF1s in different isoforms of *TUBA* gene in both human and mouse.**

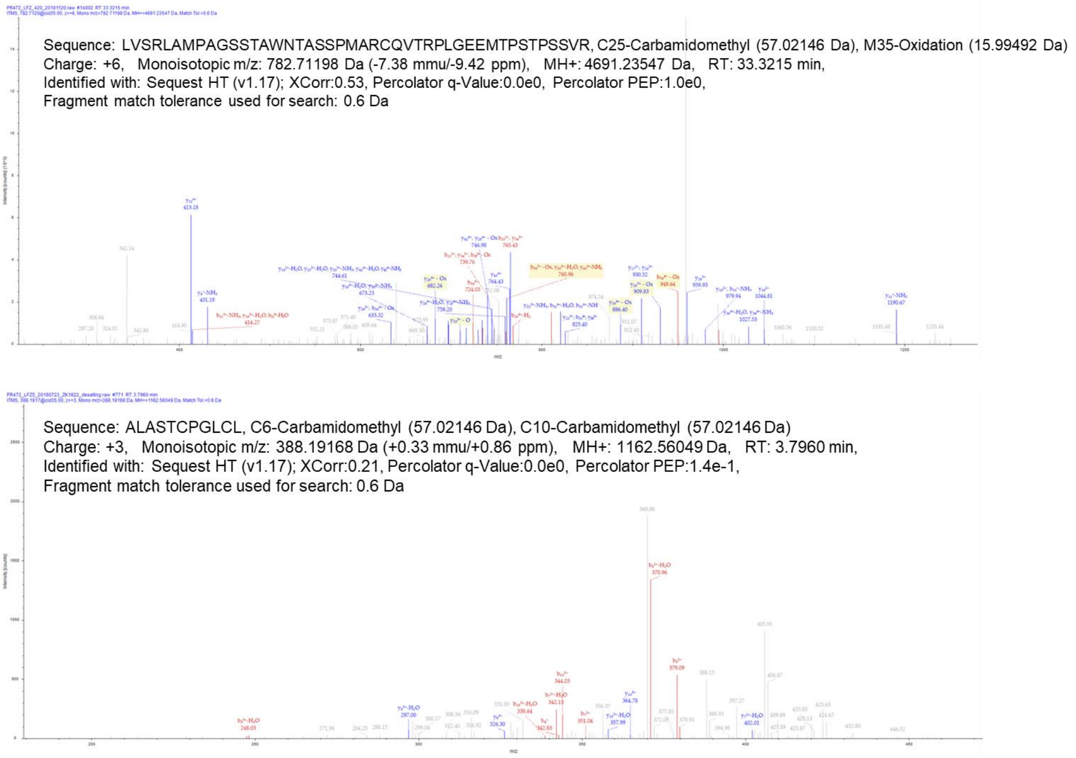


**Figure S2. Proteomic Analysis of TUBA1B-sORF1 by LC-MS/MS.**
The upper spectrum illustrates the detected peptide sequence LVSRLAMPAGSSTAWNTASSPMARCQVTRPLGEEMTPSTPSSVR and the lower spectrum depicts the peptide sequence ALASTCPGLCLC. These data provide evidence supporting the presence and post-translational modifications of TUBA1B-sORF1 peptides.

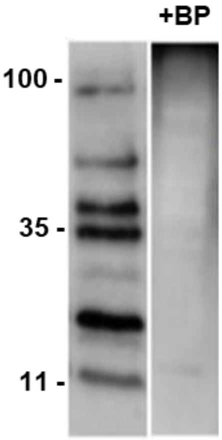


**Figure S3. Peptide blocking revealed the specificity of anti-sORF1 antibody.** HeLa cell lysates were subjected to WB with anti-sORF1 in the absence (left) and presence (right) of blocking peptide (BP).

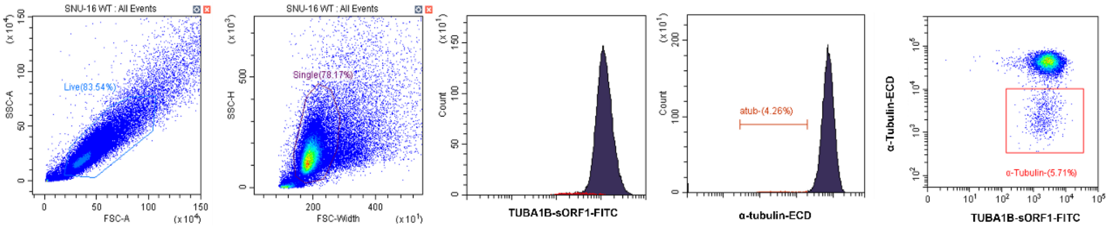


**Figure S4. Flow cytometry revealed a subpopulation in SNU-16 cells showing α-tubulin⁻/sORF1⁺.**

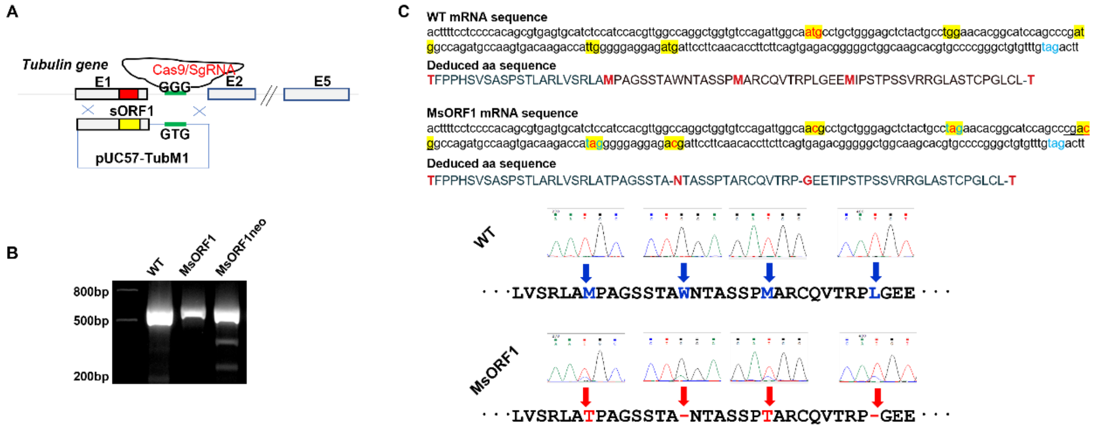


**Figure S5. Strategy to disrupt sORF1 via CRISPR/Cas9 gene editing.** (**A**) An illustration of constructs used in gene editing. (**B**) PCR plus enzyme digestion confirmed the success of gene editing. (**C**) An illustration of substituted nucleotides and deduced amino acid residue changes (upper), which was confirmed by DNA sequencing (lower).

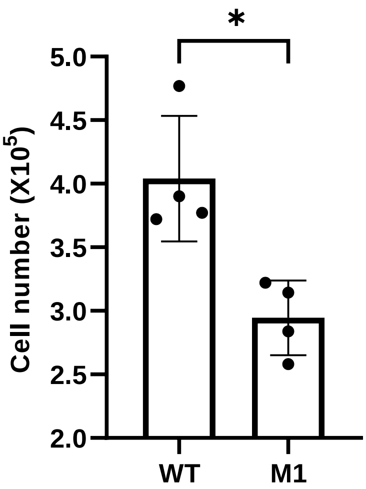


**Figure S6. Disruption of sORF1 (M1) suppressed cell proliferation as revealed by cell counting.** 5x$10^4$ cells/well were seeded in 12-well plates and cultured for 72hr with medium refreshment every other day, followed by cell counting. WT: unedited SNU-16 cells; M1: mutated SNU-16 cells. *: $p<0.05$.

| Log2 | Gene | Log2 | Gene | Log2 | Gene | Log2 | Gene | Log2 | Gene | Log2 | Gene |
|---|---|---|---|---|---|---|---|---|---|---|---|
| -4.25 | SLC43A3 | -8.28 | CTAG2 | -1.28 | PLAUR | -7.43 | LINC01153 | -1.83 | ENPP2 | 1.42 | MPP7 |
| -3.17 | ALDH3A1 | -2.31 | GJB3 | -6.07 | LIN7A | -1.11 | ADI1 | -1.02 | H3-3B | 1.56 | TMOD2 |
| -3.98 | TRIB2 | -3.41 | IL6R | -1.15 | CSRP1 | -3.75 | LY6D | -3.35 | PCAT1 | 2.54 | TRHDE-AS1 |
| -3.60 | PIWIL1 | -3.38 | ELFN2 | -3.68 | CYP27A1 | -4.66 | CAVIN3 | -5.31 | AP3B2 | 1.59 | BMP6 |
| -5.17 | GPAT2 | -3.71 | PTPN13 | -2.25 | SPNS2 | -1.71 | SMIM24 | -1.08 | FAM120C | 1.16 | PTPN14 |
| -4.41 | CLDN18 | -1.96 | POU2AF1 | -6.24 | C11orf86 | -1.52 | DIP2C | -1.79 | PLEKHO1 | 1.58 | FAM184A |
| -4.10 | TSPYL5 | -2.10 | MFGE8 | -1.30 | DMPK | -5.61 | SFMBT2 | -2.99 | ATAD3C | 1.24 | ARHGAP20 |
| -3.92 | ARHGEF9 | -4.03 | UST | -2.83 | CELF3 | -2.74 | LINC01618 | -3.41 | LRATD2 | 1.83 | OLFML2B |
| -7.21 | MAGEA12 | -2.03 | DHRS9 | -1.26 | PRSS12 | -5.51 | VEPH1 | -3.35 | CHGB | 1.52 | MEGF6 |
| -2.11 | FADS1 | -5.27 | CNTLN | -1.57 | TPPP3 | -5.54 | GPAT2P1 | -4.58 | PCSK1N | 1.84 | TRABD2B |
| -3.77 | MT1X | -5.15 | NEUROG3 | -6.22 | MAGEA6-DT | -1.79 | PLPP4 | -1.49 | CD109 | 2.50 | NR4A3 |
| -6.02 | SOHLH2 | -3.67 | WDR72 | -4.44 | PMP22 | -2.52 | BAIAP3 | -3.24 | MT1H | 2.14 | GLIPR2 |
| -2.81 | MDFI | -2.14 | RAB37 | -3.95 | SERTAD4 | -3.86 | VWA5B2 | -1.16 | LMCD1 | 1.70 | WASF3 |
| -3.65 | RAB32 | -5.11 | MS4A8 | -6.13 | OR51E1 | -1.13 | BAMBI | -1.00 | UBE2S | 1.52 | GAB2 |
| -5.26 | SLC38A5 | -2.77 | SLC7A2 | -6.00 | BTC | -1.27 | P3H4 | -1.80 | CLDN23 | 1.80 | PIGR |
| -4.35 | PPP1R14C | -2.19 | AQP3 | -1.71 | IFT27 | -6.40 | PAX4 | -5.24 | COL21A1 | 1.11 | ATP8A1 |
| -2.67 | MAGEA3 | -6.53 | FIGN | -1.55 | HOXB2 | -1.64 | COMMD7 | -1.49 | SDCBP2 | 2.12 | LINC01535 |
| -4.45 | ASCL1 | -2.00 | ARHGAP4 | -1.81 | LOXL1-AS1 | -1.22 | MAPK7 | -1.12 | SULT2B1 | 3.81 | EDNRB |
| -3.32 | CCDC88A | -2.10 | LGALS1 | -3.08 | TGFBI | -4.08 | STK33 | -5.37 | ALOX12P2 | 1.19 | CEP68 |
| -8.58 | GLI3 | -4.30 | GJA1 | -1.18 | CEACAM6 | -5.59 | KLK11 | -1.19 | PPL | 2.22 | ZBTB10 |
| -3.93 | FIRRE | -4.62 | CHST9 | -1.66 | H1-2 | -1.19 | GOLGA7B | -1.44 | ZNF785 | 1.72 | ADAM22 |
| -3.65 | LINC01561 | -3.91 | MATK | -2.17 | KRT15 | -1.44 | TMEM44-AS1 | -1.08 | KPNA2 | 1.66 | NUAK1 |
| -2.91 | RPS6KA2 | -4.26 | DMD | -6.06 | ST8SIA6-AS1 | -2.35 | GUCA2A | -3.76 | CABP7 | 1.48 | SLC1A2 |
| -2.92 | CDKN1C | -5.59 | FAM171A1 | -4.46 | HMGN5 | -4.30 | STC2 | -2.30 | CORO1A | 1.24 | STARD9 |
| -2.75 | IRS1 | -6.27 | CSAG1 | -1.38 | MB | -1.81 | TP53INP1 | -5.03 | AMOTL1 | 1.08 | PCCA |
| -4.17 | FRAS1 | -2.17 | KLK1 | -2.32 | OXTR | -2.35 | KLHL29 | -1.43 | FAM177B | 5.37 | C1QL4 |
| -2.77 | LOXL1 | -1.93 | GNG2 | -2.79 | IL1RN | -4.18 | PTGS1 | -3.16 | DNAH14 | 1.08 | TMEM51-AS1 |
| -3.12 | WNT4 | -4.45 | MIAT | -1.34 | FSIP2 | -3.59 | ZNF662 | -2.03 | SPECC1 | 1.05 | PGAP1 |
| -1.66 | HSPB6 | -1.31 | CD82 | -5.30 | SH3BGRL | -1.84 | MARCHF3 | -1.08 | POLE4 | 1.26 | CD302 |
| -7.61 | TENM1 | -2.82 | MUC5AC | -1.59 | SERTAD1 | -1.50 | H2BC12 | 1.20 | L3MBTL1 | 1.17 | RPL32P3 |
| -2.81 | TIMP1 | -6.47 | ZNF229 | -5.12 | CAMK2B | -2.11 | SLC17A4 | 2.56 | Y_RNA | 1.65 | SORBS1 |
| -2.74 | TANC2 | -6.52 | P2RX5 | -1.38 | SYT13 | -5.37 | LINC01915 | 1.32 | CDON | 1.36 | ZNF90 |
| -7.76 | LINC02864 | -5.50 | C3orf14 | -4.20 | PIP4P2 | -3.70 | CACNA1A | 1.15 | MYLK | 1.85 | SPATA18 |
| -1.91 | PPP1R2 | -6.46 | ZFPM2-AS1 | -5.83 | KLK12 | -5.63 | SSTR5 | 4.51 | MUC6 | 1.84 | RNF125 |
| -1.93 | FLRT3 | -3.36 | LUM | -1.53 | ATG4A | -3.19 | LIMCH1 | 6.09 | MME | 1.02 | RASA3 |
| -3.74 | MT1G | -2.89 | SUSD2 | -1.11 | NMI | -2.90 | TAMALIN | 4.20 | ZDHHC2 | 1.01 | FOXC1 |
| -7.64 | GTSF1 | -3.81 | CACNA1H | -5.87 | PPARGC1A | -1.56 | KATNAL2 | 2.62 | ZIC2 | 1.13 | PPM1L |
| -3.99 | CCN3 | -3.01 | CALB2 | -3.04 | FKBP10 | -1.42 | PKP1 | 5.19 | LGI2 | 1.43 | SBSPON |
| -4.40 | TPD52L1 | -4.13 | TUSC3 | -1.89 | FADS2 | -5.14 | SETBP1 | 2.76 | ATE1 | 1.35 | IQCH |
| -3.53 | RPS6KA6 | -2.16 | H2BC5 | -3.14 | MALRD1 | -1.77 | ATOH8 | 3.01 | ZIC5 | 1.22 | TBC1D2 |
| -7.51 | FAM155B | -6.10 | CHGA | -1.44 | CCDC92 | -5.63 | LRRC74B | 2.13 | LYZ | 1.13 | MUC1 |
| -2.02 | SERPINH1 | -4.96 | PRSS2 | -1.50 | MAST4 | -1.86 | CCDC149 | 3.01 | CAMKV | 1.07 | FAM160A1 |
| -7.38 | TBX5 | -3.10 | ST3GAL6 | -5.30 | CYP4F35P | -2.92 | PHLDA3 | 1.94 | GAS6 | 1.20 | ARL10 |
| -5.27 | RRAS2 | -1.27 | BST2 | -4.34 | NR0B2 | -1.59 | CRIP2 | 1.47 | CYP2W1 | 1.43 | ALDH5A1 |
| -5.61 | SORBS2 | -1.68 | LINC01278 | -2.63 | DBN1 | -5.26 | CSAG4 | 2.12 | ZYG11A | 2.88 | IGDCC4 |
| -7.48 | NEBL | -3.06 | KIF19 | -5.81 | PADI3 | -3.05 | COL7A1 | 1.62 | PRLR | 1.60 | IL27RA |
| -3.55 | NR2F1 | -1.06 | LSG1 | -2.56 | TFF1 | -1.17 | SMOX | 2.25 | SAMD4A | 1.10 | PITPNM3 |
| -4.25 | NEUROD1 | -2.26 | APOL4 | -6.37 | SCGN | -1.46 | AVEN | 1.45 | DNAJC6 | 1.01 | CHST5 |
| -6.61 | NUP210 | -1.34 | PLEKHG4 | -1.25 | PID1 | -3.32 | MYT1 | 2.82 | PPM1H | 1.70 | ETV7 |
| -2.36 | HS6ST2 | -2.87 | CIART | -1.24 | GULP1 | -1.21 | SLC44A5 | 2.07 | CDH12 | 2.53 | SYNPO |
| -1.12 | AMIGO2 | -2.96 | OGFRL1 | -4.76 | ZNF736 | -3.95 | ARX | 1.90 | THSD4 | 1.02 | WDR27 |
| -1.40 | KRT80 | -3.41 | RUSC2 | -2.03 | ALPG | -1.11 | ETV1 | 2.77 | TRHDE | 2.99 | SLX1B |
| -7.03 | FAR2P2 | -5.99 | SSTR5-AS1 | -4.83 | H19 | -1.92 | ROBO1 | 1.50 | UBE2E2 | 1.30 | GGT1 |
| -3.06 | FOXL1 | -1.15 | RASD2 | -4.79 | ZNF681 | -2.09 | IFI27 | 3.38 | PPP1R1B | 1.05 | METTL7A |
| -3.29 | ST3GAL1 | -1.88 | FFAR2 | -1.45 | PROM2 | -1.33 | H2AC6 | 1.90 | PRUNE2 | 1.16 | UBE2Q2P2 |
| -2.80 | ADGRF1 | -1.42 | TXNDC15 | -3.22 | MT2A | -1.96 | MFAP3L | 1.41 | SOAT1 | 1.69 | SLC10A5 |
| -5.13 | SCG3 | -1.19 | SH3KBP1 | -4.71 | LINC02418 | -1.06 | COPS9 | 1.11 | KCTD12 | 1.51 | MYRIP |
| -5.32 | NOVA1 | -1.13 | ALDH3A2 | -2.80 | SERPINB8 | -1.01 | SERPINB5 | 1.52 | GPA33 | 1.45 | CHDH |
| -3.42 | ST18 | -6.14 | RTL8B | -5.68 | RGS7 | -1.38 | EMP1 | 2.44 | TTYH2 | 3.96 | SNORA75 |
| -4.43 | LAMA4 | -2.25 | SLC6A20 | -1.46 | MITF | -2.07 | PRECSIT | 2.25 | LY75 | 1.02 | SEMA4G |
| -4.50 | EPHB3 | -1.11 | CYTH2 | -3.32 | BMP4 | -1.69 | DSC3 | 1.09 | RETREG1 | | |
| -4.63 | F5 | -5.65 | PRKAA2 | -4.11 | KLF2 | -1.02 | FAM50B | 1.54 | HES7 | | |
| -5.02 | RUNX3 | -2.45 | MIR22HG | -5.79 | PRSS1 | -4.19 | TRIM54 | 1.03 | METTL7B | | |
| -5.24 | APBB1IP | -4.94 | SOX21 | -4.91 | APOB | -3.66 | PALD1 | 1.35 | IL17RB | | |

**Figure S7. A full list of the genes changed by disruption of sORF1 and detected by RNA-seq as compared to unedited cells.**

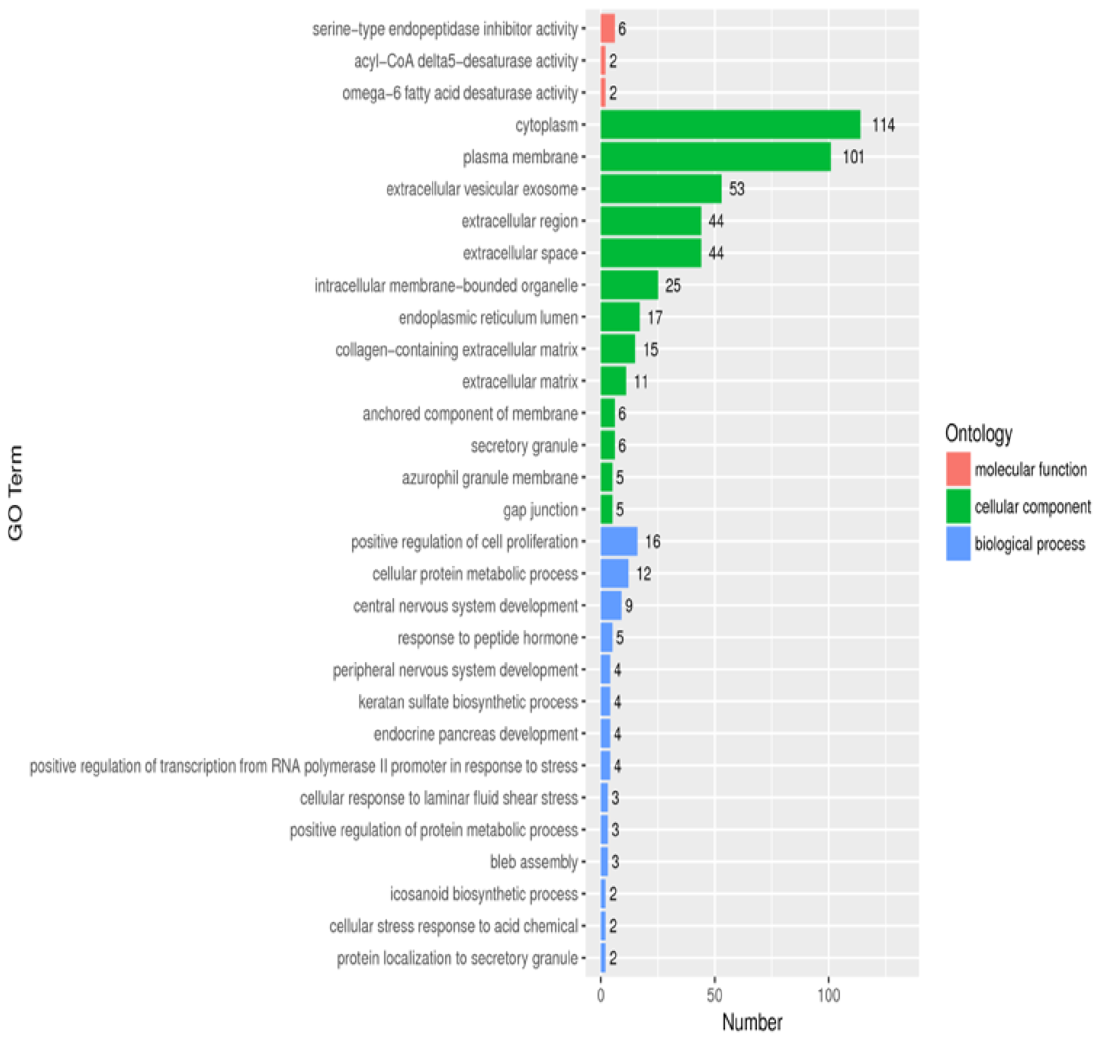


**Figure S8. Functional ontology of genes changed listed in Figure S7.**

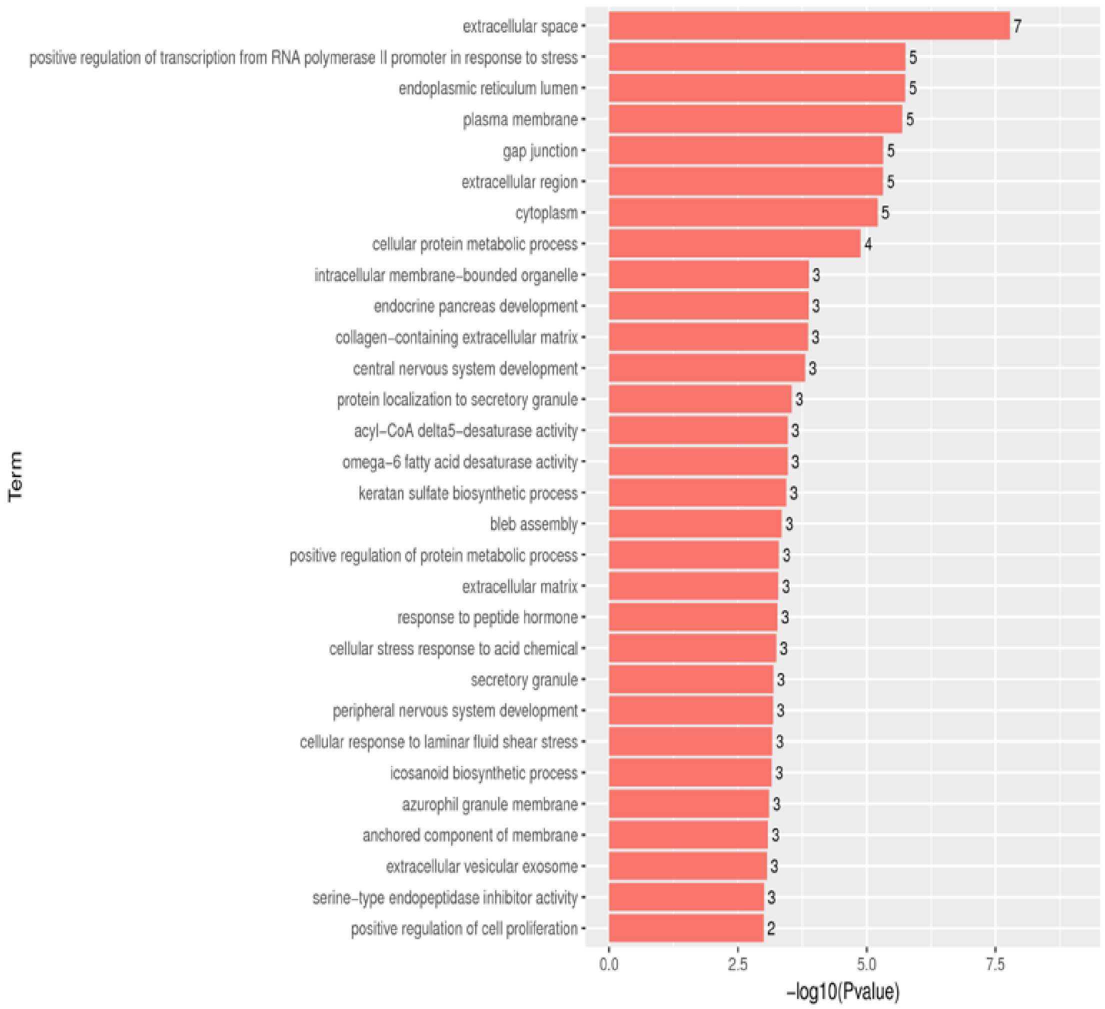


**Figure S9. Ontology of cellular processes for the genes listed in Figure S7.**

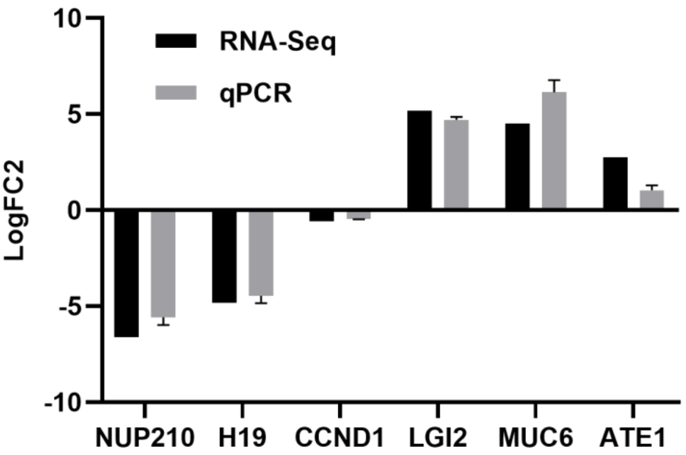


**Figure S10. Validation of significantly regulated genes identified from RNA-sequencing using RT-qPCR.**

| Acc No. | ID | MW |
|---|---|---|
| P49327 | FASN | 273 kDa |
| Q14980 | NUMA1 | 238 kDa |
| Q8WWI1 | LMO7 | 193 kDa |
| O00571 | DDX3X | 73 kDa |
| P27816 | MAP4 | 121 kDa |
| Q99623 | PHB2 | 33 kDa |
| Q13428 | TCOF1 | 152 kDa |
| Q7Z2W4 | ZC3HAV1 | 101 kDa |
| Q92945 | KHSRP | 73 kDa |
| P25705 | ATP5F1A | 60 kDa |
| P50990 | CCT8 | 60 kDa |
| Q04637 | EIF4G1 | 175 kDa |
| Q8TES7 | FBF1 | 125 kDa |
| P49207 | RPL34 | 13 kDa |
| P23588 | EIF4B | 69 kDa |
| Q99575 | POP1 | 115 kDa |
| O14974 | PPP1R12A | 115 kDa |
| Q01650 | SLC7A5 | 55 kDa |
| Q9UKV3 | ACIN1 | 152 kDa |
| Q6P2Q9 | PRPF8 | 274 kDa |
| Q68DQ2 | CRYBG3 | 331 kDa |
| Q14839 | CHD4 | 218 kDa |
| P61927 | RPL37 | 11 kDa |
| Q15233 | NONO | 54 kDa |
| Q15233 | NONO | 54 kDa |
| P23246 | SFPQ | 76 kDa |
| P46087 | NOP2 | 89 kDa |
| Q8IVT2 | MISP | 75 kDa |
| Q9NXV6 | CDKN2AIP | 61 kDa |
| P35232 | PHB | 30 kDa |
| P28331 | NDUFS1 | 79 kDa |
| P10809 | HSPD1 | 61 kDa |
| Q00341 | HDLBP | 141 kDa |
| P61978 | HNRNPK | 51 kDa |
| Q86W92 | PPFIBP1 | 114 kDa |
| Q13151 | HNRNPA0 | 31 kDa |
| Q9UHD8 | SEPTIN9 | 65 kDa |
| Q9C0C2 | TNKS1BP1 | 182 kDa |
| Q9UHB6 | LIMA1 | 85 kDa |
| Q13283 | G3BP1 | 52 kDa |
| Q92841 | DDX17 | 80 kDa |
| P48444 | ARCN1 | 57 kDa |
| Q8N684 | CPSF7 | 52 kDa |
| Q9UK61 | TASOR | 189 kDa |
| Q9H9B1 | EHMT1 | 141 kDa |
| Q9Y2D5 | AKAP2 | 95 kDa |
| O15027 | SEC16A | 252 kDa |
| Q9Y566 | SHANK1 | 225 kDa |
| P46013 | MKI67 | 359 kDa |
| P46940 | IQGAP1 | 189 kDa |
| O60763 | USO1 | 108 kDa |
| P09651 | HNRNPA1 | 39 kDa |
| P41252 | IARS1 | 145 kDa |

| Acc No. | ID | MW |
|---|---|---|
| Q15286 | RAB35 | 23 kDa |
| O60264 | SMARCA5 | 122 kDa |
| P50991 | CCT4 | 58 kDa |
| P62081 | RPS7 | 22 kDa |
| Q14152 | EIF3A | 167 kDa |
| Q6B0B8 | TIGD3 | 52 kDa |
| Q5T1M5 | FKBP15 | 134 kDa |
| Q8WXX0 | DNAH7 | 461 kDa |
| P53990 | IST1 | 40 kDa |
| Q9BXJ1 | C1QTNF1 | 32 kDa |
| P78344 | EIF4G2 | 102 kDa |
| Q06187 | BTK | 76 kDa |
| P31948 | STIP1 | 63 kDa |
| P62258 | YWHAE | 29 kDa |
| Q9BQE3 | TUBA1C | 50 kDa |
| O15541 | RNF113A | 39 kDa |
| O75717 | WDHD1 | 126 kDa |
| P08195 | SLC3A2 | 68 kDa |
| P0DN79 | CBSL | 61 kDa |
| P12270 | TPR | 267 kDa |
| P23396 | RPS3 | 27 kDa |
| P27338 | MAOB | 59 kDa |
| P43897 | TSFM | 35 kDa |
| P51991 | HNRNPA3 | 40 kDa |
| P55072 | VCP | 89 kDa |
| P55884 | EIF3B | 92 kDa |
| P60228 | EIF3E | 52 kDa |
| P60842 | EIF4A1 | 46 kDa |
| P61006 | RAB8A | 24 kDa |
| P62195 | PSMC5 | 46 kDa |
| Q08J23 | NSUN2 | 86 kDa |
| Q13200 | PSMD2 | 100 kDa |
| Q15046 | KARS1 | 68 kDa |
| Q5H9R7 | PPP6R3 | 98 kDa |
| Q5JTV8 | TOR1AIP1 | 66 kDa |
| Q5SW79 | CEP170 | 175 kDa |
| Q6WRI0 | IGSF10 | 291 kDa |
| Q7L014 | DDX46 | 117 kDa |
| Q7Z417 | NUFIP2 | 76 kDa |
| Q86UE4 | MTDH | 64 kDa |
| Q8IX12 | CCAR1 | 133 kDa |
| Q8TDD1 | DDX54 | 99 kDa |
| Q92621 | NUP205 | 228 kDa |
| Q969V3 | NCLN | 63 kDa |
| Q96KR1 | ZFR | 117 kDa |
| Q99459 | CDC5L | 92 kDa |
| Q99567 | NUP88 | 84 kDa |
| Q99729 | HNRNPAB | 36 kDa |
| Q9BTC0 | DIDO1 | 244 kDa |
| Q9H0A0 | NAT10 | 116 kDa |
| Q9H0D6 | XRN2 | 109 kDa |
| Q9UDY2 | TJP2 | 134 kDa |
| Q9Y265 | RUVBL1 | 50 kDa |

| Acc No. | ID | MW |
|---|---|---|
| Q9UHR4 | BAIAP2L1 | 57 kDa |
| P40429 | RPL13A | 24 kDa |
| Q5HYW3 | RTL5 | 65 kDa |
| Q5T8P6 | RBM26 | 114 kDa |
| Q8NE71 | ABCF1 | 96 kDa |
| Q15020 | SART3 | 110 kDa |
| P38919 | EIF4A3 | 47 kDa |
| Q9NQ55 | PPAN | 53 kDa |
| Q01518 | CAP1 | 52 kDa |
| P17655 | CAPN2 | 80 kDa |
| P06756 | ITGAV | 116 kDa |
| P00533 | EGFR | 134 kDa |
| Q8N163 | CCAR2 | 103 kDa |
| Q99856 | ARID3A | 63 kDa |
| P62306 | SNRPF | 10 kDa |
| Q92616 | GCN1 | 293 kDa |
| Q14974 | KPNB1 | 97 kDa |
| O15405 | TOX3 | 63 kDa |
| Q14534 | SQLE | 64 kDa |
| Q15637 | SF1 | 68 kDa |
| Q9H2P0 | ADNP | 124 kDa |
| Q5T5S1 | CCDC183 | 63 kDa |
| O94875 | SORBS2 | 124 kDa |
| P49756 | RBM25 | 100 kDa |
| O15117 | FYB1 | 85 kDa |
| Q9BRD0 | BUD13 | 71 kDa |
| Q99439 | CNN2 | 34 kDa |
| Q92538 | GBF1 | 206 kDa |
| Q8TEQ6 | GEMIN5 | 169 kDa |
| Q9Y262 | EIF3L | 67 kDa |
| O00400 | SLC33A1 | 61 kDa |
| Q53T59 | HS1BP3 | 43 kDa |
| P80723 | BASP1 | 23 kDa |
| Q8NDG6 | TDRD9 | 156 kDa |
| Q9P013 | CWC15 | 27 kDa |
| P78346 | RPP30 | 29 kDa |
| Q9Y4E1 | WASHC2C | 145 kDa |
| P46020 | PHKA1 | 137 kDa |
| Q969S3 | ZNF622 | 54 kDa |
| Q6ZSZ5 | ARHGEF18 | 152 kDa |
| P51532 | SMARCA4 | 185 kDa |
| P46100 | ATRX | 283 kDa |
| O60840 | CACNA1F | 221 kDa |
| Q7Z5K2 | WAPL | 133 kDa |
| Q86YP4 | GATAD2A | 68 kDa |
| P02786 | TFRC | 85 kDa |
| Q9BUJ2 | HNRNPUL1 | 96 kDa |
| Q9NX46 | ADPRS | 39 kDa |
| Q6YN16 | HSDL2 | 45 kDa |
| A8MX76 | CAPN14 | 80 kDa |
| O43395 | PRPF3 | 78 kDa |
| Q8WXG6 | MADD | 183 kDa |

**Figure S11. A list of sORF1 associated proteins identified by BioID plus proteomics analysis.**

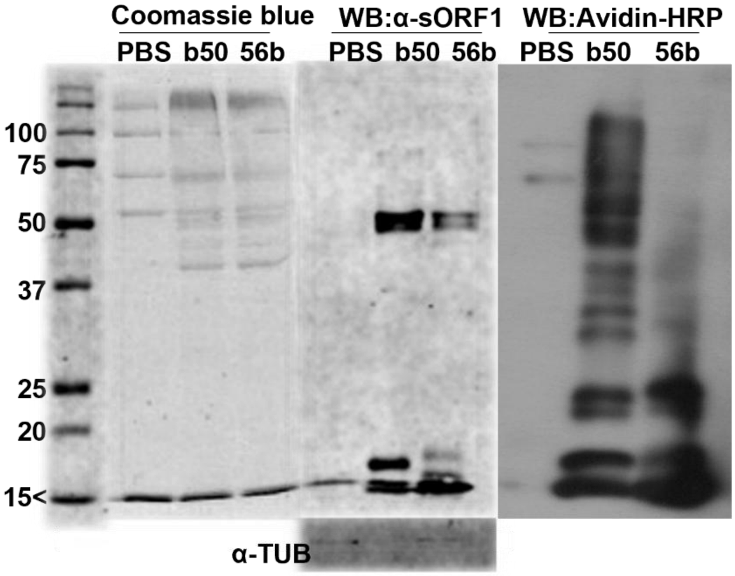


**Figure S12. The C-terminal of sORF1 is involved in sORF1-mediated post-translational modification of substrate proteins.** HeLa cell lysates (500µg) were incubated with biotinylated sORF1 (5µg) in PBS at 37ºC for 1 hour, followed by streptavidin-magnetic beads pulldown, reduced SDS-PAGE (0.2M DTT), Coomassie blue staining and WB with anti-sORF1 or Avidin-HRP. PBS was included as vehicle control. b50A: N-terminal biotinylated sORF1; 56Ab: C-terminal biotinylated sORF1 with the additional 6 amino acids (LVSRLA).
